# Full-likelihood genomic analysis clarifies a complex history of species divergence and introgression: the example of the *erato-sara* group of *Heliconius* butterflies

**DOI:** 10.1101/2021.02.10.430600

**Authors:** Yuttapong Thawornwattana, Fernando A. Seixas, Ziheng Yang, James Mallet

## Abstract

Introgressive hybridization plays a key role in adaptive evolution and species diversification in many groups of species. However, frequent hybridization and gene flow between species makes estimation of the species phylogeny and key population parameters challenging. Here, we show that by accounting for phasing and using full-likelihood analysis methods, introgression histories and population parameters can be estimated reliably from whole-genome sequence data. We employ full-likelihood methods under the multispecies coalescent (MSC) model with and without gene flow to analyze the genomic data from six members of the *erato*-*sara* clade of *Heliconius* butterflies and infer the species phylogeny and cross-species introgression events. The methods naturally accommodate random fluctuations in genealogical history across the genome due to deep coalescence. To avoid heterozygote phasing errors in haploid sequences commonly produced by genome assembly methods, we process and compile unphased diploid sequence alignments and use analytical methods to average over uncertainties in heterozygote phase resolution. There is robust evidence for introgression across the genome, both among distantly related species deep in the phylogeny and between sister species in shallow parts of the tree. We obtain chromosome-specific estimates of key population parameters such as introgression directions, times and probabilities, as well as species divergence times and population sizes for modern and ancestral species. We confirm ancestral gene flow between the *sara* clade and an ancestral population of *H. telesiphe*, a likely hybrid speciation origin for *H. hecalesia*, and gene flow between sister species *H. erato* and *H. himera*. Inferred introgression among ancestral species also explains the history of two chromosomal inversions deep in the phylogeny of the group. This study illustrates how a full-likelihood approach based on the multispecies coalescent makes it possible to extract rich historical information of species divergence and gene flow from genomic data.

## Introduction

Thanks to increasing availability of genomic data and advances in analytical methods (Sousa and Hey 2013; Payseur and Rieseberg 2016), hybridization or introgression among species has been detected in a variety of organisms, including *Anopheles* mosquitoes (Fontaine et al. 2015), *Panthera* cats (Figueiró et al. 2017) and cichlid fishes (Malinsky et al. 2018), as well as *Heliconius* butterflies (Dasmahapatra et al. 2012; Jay et al. 2018; Edelman et al. 2019; Kozak et al. 2021). Introgression is increasingly recognized to be important in the transfer of genetic diversity across species, and it likely contributes to adaptive evolution (Mallet et al. 2016; Taylor and Larson 2019; Edelman and Mallet 2021). Understanding gene flow and the role it plays during divergence is today seen as important for a fuller understanding of speciation (Pinho and Hey 2010; Feder et al. 2012).

A common approach for inferring the species phylogeny from genomic data has been to estimate phylogenetic trees based on concatenated genomic data. However, gene flow and other factors can lead to phylogenetic discordance within whole genome data. Given a species phylogeny, summary methods such as the *D* statistic (or ABBA-BABA test) (Patterson et al. 2012) are then often used to detect cross-species gene flow. This whole-genome test for gene flow is useful, but it ignores information in the local variation of the gene tree across the genome, and is unable to infer gene flow between sister species or the direction and timings of gene flow between non-sister species. More recently, phylogenetic analysis of sliding windows across the genome have been used to investigate the causes for phylogenetic discordance, particularly introgression. Sliding-window/concatenation analysis is a useful descriptive tool for exploratory analysis of genomic data (Martin and Van Belleghem 2017), but it could run into difficulties when used in inference. The assumption of a single gene tree for the whole sliding window (which may have a size of 10kb or 100kb) may be unrealistic. Furthermore, the proportions of gene trees among the sliding windows may not have a simple biological interpretation. Even under a model of complete isolation, large fluctuations in gene tree topology and branch lengths are expected due to natural coalescent fluctuations (Barton 2006): the probability that the (true) gene tree has a different topology from the species tree can range anywhere from near 0% to near 100%, depending on the parameters in the model. Estimated gene trees are affected by phylogenetic errors which are known to inflate the gene-tree discordance (Yang 2002). Introgression adds further variation. Furthermore, proportions of estimated gene trees from sliding windows are sensitive to the window size: with small windows, the results may be affected by phylogenetic errors, whereas with large windows, one or two gene trees will dominate due to the averaging effects, even though the average gene tree for the sliding window may differ from the species tree (Roch and Steel 2015). Thus it may not be straightforward to use the proportions of gene tree topologies from sliding windows to infer the presence of gene flow or to estimate its strength. For example, a previous sliding-window phylogenomic analysis of the *Anopheles gambiae* group of African mosquitoes appears to have produced an incorrect backbone species tree (Fontaine et al. 2015), as a coalescent-based re-analysis suggested a different species tree that is more consistent with chromosomal inversions and with known patterns of gene flow among the species (Thawornwattana et al. 2018).

Another common issue is the treatment of heterozygous sites in unphased diploid sequence data. A standard practice in genome assembly has been to “haploidify” the diploid sequence to a single haploid sequence for each genome, with each heterozygous site represented by only one nucleotide base, e.g., the one with more reads. This loses half of the information, and, worse still, produces chimeric haplotypes that do not exist in nature because the genotypic phase of multiple heterozygous sites is resolved effectively at random. Recent simulation studies (Andermann et al. 2019; Huang et al. 2021) found that such haploid consensus sequences can lead to serious biases in downstream phylogenomic analyses if the species tree is shallow, with species divergence times comparable to coalescent times. The impact of phasing errors in haploidified sequences on estimation of species trees and inference of gene flow is poorly understood. The BPP program (Yang 2015; Flouri et al. 2018) implements the algorithm of Gronau et al. (2011), which enumerates and averages over all possible phase resolutions of heterozygote sites, weighting them according to their likelihood based on the sequence alignment at the locus. In simulations, this approach performed nearly as well as analysis of fully phased diploid genomes (which could be generated, for instance, by costly single-molecule cloning and sequencing) (Huang et al. 2021).

Here, we test coalescent-based full-likelihood phylogenomic approaches that explicitly account for deep coalescence and introgression as sources of genealogical variation across the genome while accounting for unphased diploid sequences probabilistically. One approach is based on a multispecies coalescent model with introgression (MSci) (Wen and Nakhleh 2018; Zhang et al. 2018), implemented in the program BPP (Flouri et al., 2020). In this approach, introgression is modelled as discrete events that occur at particular time points in the past. Another approach is based on an isolation-with-migration (IM) model (Hey and Nielsen 2004; Hey 2010) implemented in the program 3s (Zhu and Yang 2012; Dalquen et al. 2017), which allows for continuous migration at a constant rate per generation between two species after their divergence. Advantages of full-likelihood methods over approximate coalescent methods or summary statistics include making full use of information in the sequence data and properly accounting for uncertainty in gene trees (Xu and Yang 2016; Jiao et al. 2021). These methods allow us not only to infer the presence of gene flow, but also to estimate its direction, timing and magnitude, along with species divergence times and effective population sizes. Estimation of these key evolutionary parameters from genome-scale sequence data can provide powerful insights into the divergence history of species, and a basis for further investigations of the evolution of adaptive traits of interest.

We use *Heliconius* butterflies as a trial group to explore the power of these methods. *Heliconius* is a rapidly radiating group in tropical America (Kozak et al. 2015). They are unpalatable to predators and are perhaps best known for mimicry, which causes multiple unrelated sympatric species to converge on similar wing patterns in local regions as a common warning sign to deter predators (Bates 1862; Müller 1879). The genus *Heliconius* comprises two major clades, the *erato-sara* clade and the *melpomene*-silvaniform clade, which diverged around 10–12 million years ago (Ma) in the Miocene (Kozak et al. 2015). Natural hybridization among species is well-documented within each clade (Mallet et al. 2007). The prevalence of introgression between species coupled with rapid radiation of species makes estimation of the species phylogeny challenging (Kozak et al. 2021). As a result, our understanding of the history of species divergence and introgression in *Heliconius* remains incomplete. Here, we estimate the species phylogeny and introgression history of six species in the *erato-sara* clade of *Heliconius* butterflies from whole-genome sequence data.

Most previous studies of the genus *Heliconius*, including the *erato* group, have focused on evolutionary relationships and gene flow at specific regions of the genome, especially the colour pattern loci responsible for phenotypic variation in the mimetic wing patterns, typically in a few species, and mainly in the *melpomene*-silvaniform clade where gene flow appears to be more prevalent (Dasmahapatra et al. 2012; Nadeau et al. 2013; Martin and Van Belleghem 2017; Jay et al. 2018; Balaban et al. 2019). Other studies have focused on wing-colour mimicry loci between two species with comimic races, *H. erato* (*erato-sara* clade) and *H. melpomene* (*melpomene*-silvaniform clade) (Hines et al. 2011; Reed et al. 2011).

Earlier molecular phylogenetic studies of *Heliconius* were based on a small number of loci (Brower 1994; Brower and Egan 1997; Beltrán et al. 2002; Beltrán et al. 2007; Kozak et al. 2015), revealing variation in gene genealogies among loci. In particular, Kozak et al. (2015) employed BUCKy (Larget et al. 2010) and *BEAST (Heled and Drummond 2010) to account for the heterogeneity of gene genealogies across loci. Hybridization and introgression were acknowledged but not directly accounted for in the analysis. Kozak et al. (2021) analyzed genome-wide coding loci from >100 individuals from 40 *Heliconius* species to estimate the species tree using approximate multispecies coalescent (MSC) methods such as ASTRAL (Mirarab et al. 2014) and MP-EST (Liu et al. 2010), and to test for introgression using a range of summary methods including the *D* statistic (or ABBA-BABA test) (Patterson et al. 2012), *f*_4_(Reich et al. 2009), and an approximate multispecies coalescent with introgression (MSci) method PhyloNet/MPL (Yu and Nakhleh 2015). However, *D* and *f*_4_ statistics use site-pattern counts averaged over the genome, ignoring information in genealogical fluctuations, while PhyloNet/MPL takes estimated gene trees as input without accounting for phylogenetic reconstruction errors and uncertainties. Van Belleghem et al. (2017) estimated the species phylogeny of the *erato* clade from concatenated autosomal SNP data, focusing on wing colour pattern loci among different geographic races of *H. erato*.

Recently, Edelman et al. (2019) conducted phylogenetic analyses with sixteen new genome assemblies of *Heliconius* species using ASTRAL to analyze gene trees and from sliding windows (Mirarab et al. 2014). Introgression was tested using *D* statistics (Patterson et al. 2012) and a new test called QuIBL based on internal branch lengths in estimated triplet gene trees. Summary methods such as *D* statistics are based on genome-wide averages and ignore information from genealogical variation across the genome in multilocus sequence data. QuIBL may be affected by sampling errors in estimated gene trees and branch lengths. Those methods typically cannot infer gene flow between sister species and have limited ability to characterize the direction, timing, and magnitude of gene flow (as measured by the migration rate or introgression probability) between non-sister species. In Edelman et al. (2019), MSci network models were inferred using PhyloNet/MCMC_SEQ (Wen and Nakhleh 2018), a Bayesian Markov chain Monte Carlo (MCMC) method based on sequence alignments. However, the method is computationally applicable only to a small number of loci, leading to considerable uncertainty of species tree topology and timing and directions of introgression (Edelman et al. 2019). Finally, Massardo et al. (2020) re-analyzed genome data from the *erato-sara* clade after adding two additional genomes using approaches similar to those of Edelman et al. (2019). Given known limitations of concatenation, the sliding-windows, and summary methods, we hypothesized that our methodology could add additional resolution using the genomic data employed by Edelman et al. (2019).

In this paper, we processed the raw reads for the genomic data of Edelman et al. (2019) for six species in the *erato-sara* clade to compile alignments of unphased diploid genomic sequences, and used them as input data for analysis using BPP.

## Methods

### Genome sequence data and genotyping

We used the raw sequencing read data from six *Heliconius* species in the *erato-sara* clade and *H. melpomene* (as an outgroup) generated by Edelman et al. (2019) (**Table S1**). Sequencing reads were filtered for Illumina adapters using cutadapt v1.8.1 (Martin, 2011) and then mapped to the chromosome-level genome assembly of *H. erato demophoon* v1 (Van Belleghem et al. 2017) available from lepbase.org, using BWA mem v0.7.15 (Li 2013) with default parameters and marking short split hits as secondary. Mapped reads were sorted and duplicate reads removed using sambamba v0.6.8 (Tarasov et al. 2015). Realignment around indels was performed with the Genome Analysis Toolkit (GATK) v3.8 *RealignerTargetCreator* and *IndelRealigner* modules (McKenna et al. 2010; DePristo et al. 2011) in order to reduce the number of indel miscalls. Read depth and other relevant read alignment quality control metrics were computed using QualiMap v2.2.1 (Okonechnikov et al. 2016).

Genotype calling was performed on each individual separately with bcftools v1.5 (Li et al. 2009) *mpileup* and *call* modules (Li 2011), using the multiallelic-caller model (*call -m*) and requiring a minimum base and mapping quality of 20. Genotype calls were filtered using the bcftools *filter* module. Both invariant and variant sites were required to have a minimum quality score (QUAL) of 20. Furthermore, we required that each genotype had a genotype quality score (GQ) ≥20 and a read depth (DP) satisfying max(1/2*meanDP, 20) ≤ DP ≤ 2*meanDP where meanDP is the average read depth of the sample. The meanDP-based filters were used to reduce erroneous calls when the read depths were too low or too high from the sample average, accounting for variation of sequencing depths across individuals. The DP ≥ 20 filter was chosen to minimize genotyping error rates (see next section) while retaining a sufficiently large number of loci across the genome. At our estimated base-calling error (∼0.1%), this filter achieved the genotype-calling error of <0.05% (see next section) (**Fig. 1, Tables S2–S3**). For the female Z chromosome, we used DP ≥ 10 instead since only one chromosome copy was present. All genotypes that did not fulfill these requirements or were located within 5-bp of an indel were recoded as missing data.

**Figure 1.**
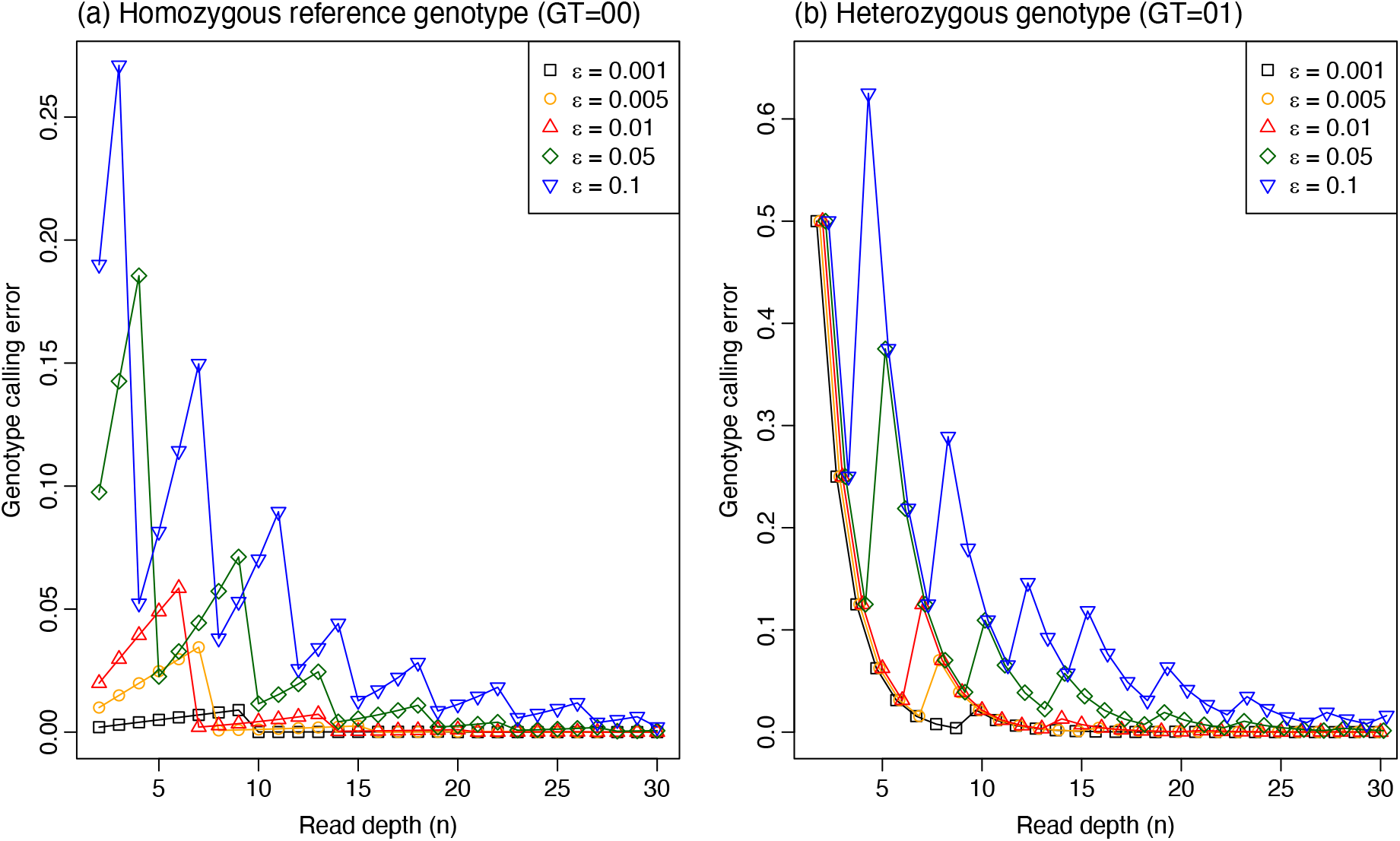
The genotype-calling error rate for a given base-calling error (*ε*) and read depth (*n*) when the true genotype is (**a**) a homozygote for the reference allele (GT = 00) or (**b**) a heterozygote (GT = 01). Note that the genotype-calling error does not decrease monotonically with the increase in *n* when *ε* is fixed, or with the reduction in *ε* when *n* is fixed. The calculation is performed using a C program written by Z.Y. that implements the ML method of Li (2011) and calculates its error rate.

### Analysis of base-calling and genotype-calling error rates to guide data compilation

The base-calling error rate was estimated from the proportion of non-matching bases in homozygous genotype calls from reads mapped to the *H. melpomene* reference genome, which was more complete than the *H. erato demophoon* reference. Only positions with homozygous genotype calls with read depth (DP) ≥ 50 were retained. Because mapping errors can bias our estimates of base-calling error rate, sites overlapping repetitive regions were excluded. For sites passing those filters, we recorded the read depth (DP) and the number of reads supporting each of the reported alleles (AD), with the difference to be the number of erroneous base calls. The base-calling error was calculated as the ratio of the number of erroneous base calls over the read depth, both summed across all sites passing filters. The calculation was done for homozygous reference genotype (GT = 00) and homozygous alternative genotype (GT = 11).

We then analyzed the genotype-calling error rate to determine a suitable cutoff for the read depth, following the maximum likelihood (ML) method for genotype-calling of Li (2011). The results were used to guide our choice of the filter (DP ≥ 20), to achieve a genotype-calling error of <0.05%.

Given the base-calling error rate (*ε*) and the read depth (*n*), the genotype-calling error rate (*e*) can be calculated by following the ML procedure of genotype calling of Li (2011). Here we assume that *ε* is the same among the reads and is independent of the true base. Given the data of *k* 1s and (*n* – *k*) 0s among the *n* reads, where 0 refers to the reference allele and 1 refers to the alternative allele, the likelihoods for the three genotypes (GT = 00, 01, and 11) are given by the binomial probabilities as

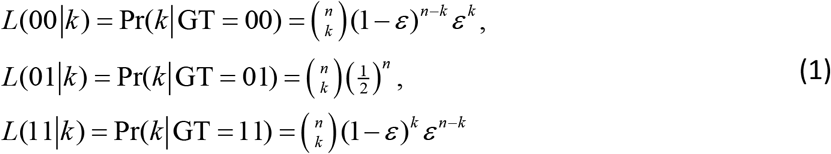

(Li 2011). The genotype achieving the highest likelihood is the called (inferred) genotype. The genotype-calling error is an average over the possible read outcomes (i.e., over *k* = 0, …, *n*)

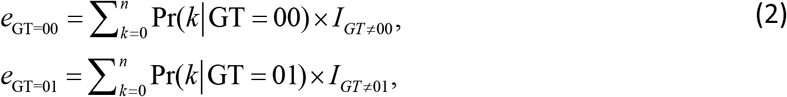

where the indicator *I*_*GT ≠*00_ is 1 if the called genotype (from data *k*) is not 00 (in error) and 0 otherwise, and *I*_*GT ≠*01_ is defined similarly.

### Multilocus datasets for BPP and 3S analyses

We prepared two sets of data, one for BPP analyses (under the MSC and MSci models) and the other for 3S analysis (under the IM model). The ‘BPP dataset’ has six species in the *erato-sara* clade. The ‘3S dataset’ has an extra species, *H. melpomene*, as an outgroup (**Table S1**) (Edelman et al. 2019). Both datasets were prepared in the same way. For each dataset, we defined coding and noncoding regions based on the gene annotation of *H. erato demophoon* v1 reference assembly (Van Belleghem et al. 2017), where coding regions included exons while noncoding regions included introns and intergenic regions. For the noncoding regions, we extracted the genotype calls for small genomic segments, referred to as loci, with sizes between 100 and 2,000 bp, and with the further requirement that any two consecutive loci must be at least 2,000 bp apart. Although linkage disequilibria largely disappear over ∼10 kb (Dasmahapatra et al. 2012), we used this minimum distance of 2 kb to retain more information in the data for species tree estimation. For each locus, we produced sequence alignments from the genotype calls, with heterozygotes represented using IUPAC codes. We applied the following filtering steps: (1) Sites within repetitive regions (based on the repeat annotation of *H. erato demophoon* v1 reference assembly, available at http://download.lepbase.org/v4/repeatmasker/) were masked; (2) loci with >50% missing data in the whole alignment were excluded; (3) for the remaining loci, sites with missing data were excluded; (4) finally, loci with ≤10 sites remaining after processing were excluded. For the coding regions, the same filtering process was applied except that there was no maximum locus size. There were 74,999 coding loci (median length of 165 and median informative sites of 3) and 92,966 noncoding loci (median length of 237, median number of informative sites of 5).

### Inferring species divergence history across the genome using multispecies coalescent model

We inferred species trees using Bayesian inference under the MSC model without gene flow implemented in BPP v4.3.0 (Yang and Rannala 2014; Rannala and Yang 2017; Flouri et al. 2018). This analysis under the MSC accounts for deep coalescence and the resulting gene tree heterogeneity along the genome but assumes no gene flow. We analyzed autosomes and the Z chromosome separately and coding and noncoding regions of autosomes separately. Two inversion regions in chromosomes 2 and 15 were analyzed separately, denoted 2b and 15b, respectively. The 2b region included part of the Herato0211 scaffold (from position 1434133) and the whole scaffolds Herato0212, Herato0213 and Herato0215 (a total of 1.95 Mb). The 15b region corresponded to Herato1505:1977997–2558395 (580 kb), which included the *cortex* gene (Herato1505:2074108– 2087841, ∼13.7 kb). Thus, there were 25 chromosomal regions in total (for 21 chromosomes, with chromosomes 2 and 15 split into three regions).

For each of the datasets (coding and non-coding loci from the Z chromosome and autosomes), we formed multilocus blocks of 100 loci and inferred the species tree for the block (**Table S4**). This block size was large enough for the inferred species tree to achieve reasonably high posterior support and yet small enough to allow for local introgression history to be reflected in the inferred ‘species tree’ for that region. Any final blocks for each chromosomal region with fewer than 40 loci were discarded due to limited information. To assess the impact of block size, we repeated the analysis using blocks of 200 loci. While the neutral coalescent process may be expected to be largely homogeneous across the genome, the introgression rate is expected to be highly variable, due to selective removal of introgressed alleles, affected by the strength of selection, local recombination rate, etc. The analysis using blocks of loci can thus capture the variation across the genome due to differential rates of gene flow. Note that both our block-based analysis and the sliding-windows analysis of Edelman et al. (2019) may capture the variation in the rate of gene flow across the genome, but there is an important difference. The 100 or 200 loci in the same block are assumed to have independent genealogical histories due to coalescent fluctuations even though they have the same underlying species tree. In the sliding-windows/concatenation analysis, all sites in the whole sliding window are assumed to share the same gene tree and divergence times. In other words, the block-based analysis assumes no recombination within each locus (100-2000 bp) while the sliding-windows analysis assumes no recombination in the window of 10kb or 50kb.

The MSC model involves two sets of parameters: the species divergence times or node ages on the species tree (*τ*) and the effective population sizes (*θ* = 4*Nμ*). Both are scaled by mutations and measured in the expected number of substitutions per site. We assigned diffuse inverse-gamma priors to those parameters, with a shape parameter of 3 and with the means to be close to rough estimates from the data in preliminary runs. Specifically, we used *θ* ∼ InvG(3, 0.04), with mean 0.04/(3 – 1) = 0.02 for all populations, the root age *τ*_0_∼ InvG(3, 0.06) for the coding loci and *τ*_0_∼ InvG(3, 0.12) for the noncoding loci. Given *τ*_0_, divergence times at other nodes of the species tree were generated from a uniform-Dirichlet distribution (Yang and Rannala 2010, eq. 2). Population-size parameters (*θ*) are integrated out analytically to improve mixing of the MCMC. The MCMC was run for 2×10^6^ iterations after a burn-in of 10^5^ iterations, with samples taken every 20 iterations. The same analysis of each block was repeated ten times using different starting species trees, and consistency among runs was used to assess convergence of the MCMC. Non-convergent runs were discarded. Samples were then combined to produce the posterior summary such as the maximum a posteriori (MAP) tree.

### Exploring gene flow scenarios using the IM model for species triplets

In order to formulate hypotheses about gene flow between species, we attempted to use several heuristic methods, including PhyloNet/MCMC_SEQ using sequences (Wen and Nakhleh 2018), PhyloNet/ML (Yu et al. 2014), and SNaQ in PhyloNetworks (Solís-Lemus et al. 2017), the latter two using estimated gene trees as input data. Our attempts were not successful. Different runs of those programs inferred different network models, which in general did not appear to be reliable, with apparently spurious introgression events around the root of the species tree.

Instead we decided to use the maximum likelihood program 3S (Dalquen et al. 2017) to estimate migration rates between all pairs of species. This implements the IM model with continuous gene flow for three species *S*_1_, *S*_2_ and *S*_3_, assuming the species tree ((*S*_1_, *S*_2_), *S*_3_), with gene flow allowed only between *S*_1_ and *S*_2_. The third species (*S*_3_) is used as an outgroup to improve parameter estimation and the power of the test (Dalquen et al. 2017). This method accommodates both deep coalescence and gene flow. While limited to three species and three sequences per locus, it can be applied to large datasets with >10,000 loci. *H. melpomene* was used as an outgroup for all pairs. The outgroup species was added to the dataset by mapping it to the *H. erato demophoon* reference genome.

We analyzed coding and noncoding regions, as well as the autosomal loci and the Z chromosome separately. However, in the 3s analysis we treated all loci in each chromosomal region as one dataset (rather than breaking them into blocks of 100 or 200 loci) (**Table S4**). We also performed an analysis that used all autosomal loci (17,428 noncoding or 28,204 coding loci; **Table S4**). Since the sequences were unphased and 3S requires phased haploid sequences, we first phased the loci using PHASE v2.1.1 (Stephens et al. 2001), resulting in two phased haploid sequences for each locus per individual. In the simulations of Huang et al. (2021), which examined the Bayesian method BPP, this approach of computational phasing produced very small biases, and the same may be expected to apply to the ML method implemented in 3S. The 3S program uses three sequences but allows multiple sequences per species. For each locus, we sampled three sequences of configurations 123, 113 and 223 with probabilities 0.5, 0.25 and 0.25, respectively. Here, 123 means one sequence from each species, etc.

Each dataset was analyzed using 3S to fit two models: MSC without migration (M0) and MSC with migration (M2; isolation-with-migration or IM). Model M0 involves two divergence time parameters (*τ*_1_ and *τ*_0_) and four effective population sizes (*θ*_1_ and *θ*_2_ for species *S*_1_ and *S*_2_, *θ*_4_ for the root, and *θ*_5_ for the ancestor of *S*_1_ and *S*_2_), while M2 involves in addition two migration rates in both directions *M*_12_ and *M*_21_. Here *M*_12_= *m*_12_*N*_2_ is the expected number of migrants from species *S*_1_ to *S*_2_ per generation, where *m*_12_ is the proportion of migrants from *S*_1_ in species *S*_2_. We use the real-world forward-in-time view to define migration parameters. Each ML analysis was repeated ten times, and the run with the highest log-likelihood value was used. The two models (M0 and M2) were compared using a likelihood ratio test (LRT), with 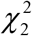 used as the null distribution. Only gene flow scenarios that passed the LRT at the 1% level were considered later.

Evidence for pairwise gene flow from the 3S analysis was used together with the inferred species trees from the blockwise BPP MSC analysis to generate an MSci model or a species tree with introgression events. There were two difficulties in this approach. First, IM (3S) and MSci (BPP) are two extreme versions of models of gene flow and either may be too simplistic to fit the real data. The IM model assumes continuous gene flow between species *S*_1_ and *S*_2_ since their split, and may be unrealistic for many of the species pairs, for example, if gene flow occurred immediately after species split but then stopped as the two species became more diverged. This is particularly the case as a branch in the 3-species tree used in the 3S analysis may represent multiple branches in the full tree for all six species. The MSci model instead assumes episodic introgression events at specific time points. Our expectation is that if introgression is episodic, the LRT based on the continuous IM model will still detect gene flow, but with distorted parameter estimates. The opposite may be true as well: if gene flow is continuous as in the IM model, fitting the MSci model will give distorted parameter estimates (Jiao et al. 2020). Second, the LRT suggested significant evidence for gene flow between most pairs of species. We prioritized introgression events that could reconcile different species trees across the genome from the above blockwise BPP analysis under the MSC model. We also took a parsimonious approach to minimize the number of introgression events by assuming gene flow only if the model of no gene flow is rejected by data: if 3s suggested gene flow between a species A and most descendant species of a node B, we assumed introgression edges between the ancestral populations.

### Estimation of parameters for species divergence and cross-species introgression

Given the MSci model constructed above, we ran BPP v4.3.0 (Flouri et al. 2020) to estimate the parameters: species divergence/introgression times (*τ*), population sizes (*θ*), and introgression probabilities (*φ*). The introgression probability is the proportion of contribution from the parental species to the hybridizing population. When we trace the genealogy of a sampled sequences backwards in time and reach a hybridization node, the sequence takes the paths of the two parental species with probabilities *φ* and 1 – *φ*, respectively. We assigned *θ* ∼ InvG(3, 0.01), with mean 0.005, *τ*_0_∼ InvG(3, 0.04) and *φ* ∼ beta(4, 2), with mean 0.75. Initial values of *φ* were set to 0.8 or 0.9. These prior settings were based on rough estimates from preliminary runs. We also considered alternative priors for *θ* (e.g. gamma distributions) and *φ* (e.g. beta(1,1)) to assess the stability of the posterior estimates when mixing and convergence were of concern. We performed inference for each of the 25 chromosomal regions, using either all coding or noncoding loci in each region (**Table S4**).

The MCMC was run for 10^6^ iterations, sampling every 100 iterations, after a burn-in of 10^6^ iterations. Ten independent runs were performed and convergence was assessed by examining consistency between runs. Non-convergent runs were discarded. For models with bidirectional introgressions (BDIs) which cause label-switching unidentifiability issues (Flouri et al. 2020), MCMC samples were post-processed before being combined for posterior summaries.

## Results

### Analysis of base-calling and genotype-calling error rates

We processed the raw reads from the whole-genome sequence data of Edelman et al. (2019) for six *Heliconius* species from the *erato-sara* clade: *H. erato demophoon, H. himera, H. hecalesia formosu*s, *H. telesiphe telesiphe, H. demeter* and *H. sara magdalena* (**Table S1**). To guide our compilation of unphased loci, we estimated the base-calling error rate in the genomic data and analyzed the genotype-calling error. The base-calling error was calculated to be ∼0.08% for the homozygous reference allele and ∼0.20% for the homozygous alternative allele, with variation among individual genomes (**Tables S2**–**S3**).

Given a base-calling error rate (*ε*), we calculated the genotype-calling error rate (*e*) for the homozygous reference genotype (GT = 00) and the heterozygous genotype (GT = 01) by using equation (2) (**Fig. 1**). Note that even with a very low base-calling error rate, the genotype-calling error can be very high, especially at low read depth. Furthermore, the genotype-calling error rate for heterozygotes is much higher than for homozygotes.

Given the read depth *n*, the genotype-calling error does not necessarily decrease when *ε* decreases. For example, when *n* = 7, the genotype-calling error for a homozygote (GT = 00) is 0.0020 at *ε* = 0.01, but rises to 0.0345 when *ε* is reduced to 0.005 (**Fig. 1**). This is due to the discrete nature of the read outcome. At *ε* = 0.01, the called genotype is 00 (the truth) when *k* = 0 or 1, is 01 when *k* = 2–5 and is 11 when *k* = 6 or 7, and the genotype-calling error is a sum over *k* = 2, …, 7 (i.e., six out of the eight cases are in error). In contrast, at *ε* = 0.005, the called genotype is 00 when *k* = 0, is 01 when *k* = 1–6 and is 11 when *k* = 7, so that the genotype-calling error is a sum over *k* = 1, …, 7 (i.e., seven out of the eight cases are in error), and is higher than at *ε* = 0.01. Similarly given *ε*, the genotype-calling error does not necessarily decrease with the increase of the read depth *n* (**Fig. 1**). For example, when *ε* = 0.01 the genotype-calling error for a homozygote is 0.0199 at *n* = 2, but rises to 0.0585 when *n* increases from 2 to 6; for those values of *n*, genotype-calling error is a sum over *k* = 1, …, *n*. Then when *n* = 7, error drops to 0.00269 as the error is a sum over *k* = 2, …, *n*. These anomalies and the strong periodicity in the genotype-calling error are both due to the discrete nature of the problem.

With the base-calling error estimated at *ε ≈* 0.001, we estimate the genotype-calling error rate to be 1.1×10^−6^ for homozygotes and 4.0×10^−4^ for heterozygotes at the read depth *n* = 20 (**Fig. 1**). We therefore filtered our data for coverage *n* ≥ 20. Since the read coverage was >60x (Edelman et al. 2019), this caused relatively little dropout.

### Gene flow variation across the genome

We first used BPP to infer species trees under the MSC model using multilocus blocks of 100 or 200 independent loci along the genome. There were 749 coding blocks and 933 noncoding blocks over the 25 chromosomal regions (21 chromosomes with chromosomes 2 and 15 each split into three regions by inversion) (**Table S4**). This is the BPP A01 analysis of Yang (2015), which explicitly accounts for deep coalescence but ignores gene flow. Ten species trees were the best estimate (i.e., the maximum *a posteriori* probability tree or MAP tree) in at least one block (**Fig. 2, Tables S5–S7**). These are referred to as trees i-x. Tree xi appeared as one of the top eight trees in the sliding-window analysis of Edelman et al. (2019) but not in our analysis. Trees i and ii accounted for over 95% of the blocks. These two trees differ only in the position of *H. hecalesia* within the *erato* clade: in tree i, *H. hecalesia* is sister to *H. telesiphe* while in tree ii, it is sister to the clade (*H. erato, H. himera*).

**Figure 2.**
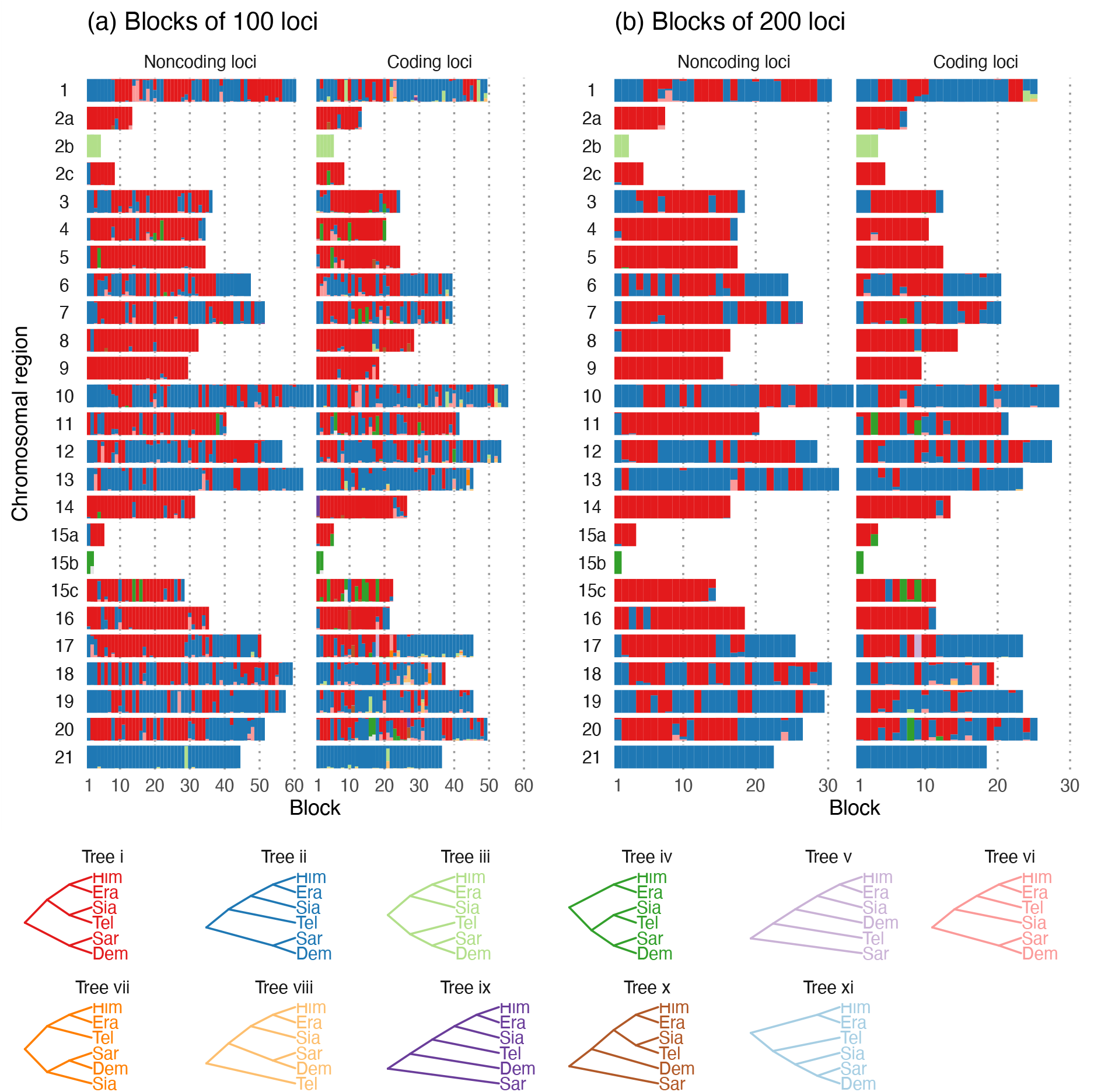
Posterior probabilities of species trees from BPP analysis of (**a**) blocks of 100 loci and (**b**) blocks of 200 loci across the genome (**Table S4**) under the MSC model without gene flow. The y-axis represents the posterior probability and ranges from 0 to 1. Trees i–x in the legend are MAP trees in at least one block. Tree colours match those in Edelman et al. (2019) (their Fig. 2). Inversion regions 2b and 15b on chromosomes 2 and 15 were analyzed separately. Era: *H. erato*, Him: *H. himera*, Sia: *H. hecalesia*, Tel: *H. telesiphe*, Dem: *H. demeter*, Sar: *H. sara*.

The species trees estimated in this analysis are expected to reflect the history of species divergences as well as cross-species introgression, and the major cause of the differences among the blocks is the variable rate of introgression along the genome. The use of 100 or 200 loci in each block helps to filter out local stochastic fluctuations in the coalescent process. The results were highly consistent between the two choices of block size and between coding and noncoding loci (**Fig. 2**), providing evidence that the patterns revealed are real rather than a result of analytical artefacts.

There was some variation in the estimated species trees across the genomic regions. Three regions in particular had species tree distributions that differed from the rest of the genome: chromosome 21 (Z chromosome), and the inversions on chromosomes 2 and 15 (2b and 15b). Chromosome 21 was the only chromosome for which tree ii is a MAP tree for almost all blocks (**Fig. 2, Table S5**). Among the autosomal regions excluding inversions, tree ii was the MAP tree in ∼40% and 46% of noncoding and coding blocks, respectively, while the corresponding proportions for tree i were 58% and 47% (**Table S5**). Thus, tree i was the autosomal majority tree. Inversion 2b had an unusual history in which *H. telesiphe* is more closely related to the *sara* clade (tree iii). In this inversion region, *H. erato, H. himera* and *H. hecalesia* share a derived inverted rearrangement relative to *H. melpomene, H. sara* and *H. demeter* (Van Belleghem et al. 2017; Davey et al. 2017; Edelman et al. 2019), consistent with trees ii and iii where these species are clustered together. Inversion 15b supported tree iv in both coding and noncoding regions, in which the (*H. telesiphe, H. hecalesia*) clade was sister to the *sara* clade instead of other members of the *erato* clade as in tree i or ii. This grouping strongly suggests that *H. telesiphe, H. hecalesia, H. demeter* and *H. sara* share the derived inverted rearrangement of this region relative to *H. erato* (Edelman et al. 2019). The 15b inversion contains the *cortex* gene that controls mimetic wing colour patterning across *Heliconius* species (Nadeau et al. 2016; Van Belleghem et al. 2017). Tree iv also appeared as a MAP tree sporadically in other parts of the autosomes, sometimes with high posterior probabilities (**Fig. 2, Table S6**). These include regions on chromosome 15 outside the inversion as well as on chromosomes 4, 5 and 11.

### Pairwise gene flow rates

Gene flow among species could reconcile different species trees from the blockwise BPP MSC analysis reported above. We investigated this by explicitly estimating gene flow rates between each pair of *erato-sara* group species using the program 3S (Zhu and Yang 2012; Dalquen et al. 2017). We used *H. melpomene* as an outgroup in all triplets for parameter estimation.

There was evidence of bidirectional gene flow between *H. telesiphe* and *H. hecalesia*, consistent with a scenario in which tree i (autosome-majority tree) and tree ii (Z chromosome tree) are related through introgression between the two species (**Fig. 3**). However, gene flow between these species was detected only for autosomal loci (both coding and noncoding) and not on the Z chromosome (**Fig. 3**). The results suggest that the prevailing topology on the Z chromosome, tree ii rather than tree i, is the true species tree. In particular, we found no evidence of gene flow in 3s analyses from *H. erato* and *H. himera* towards *H. hecalesia* (**Fig. 3**), which is necessary to explain tree ii as a result of introgression if tree i were the true species tree.

**Figure 3.**
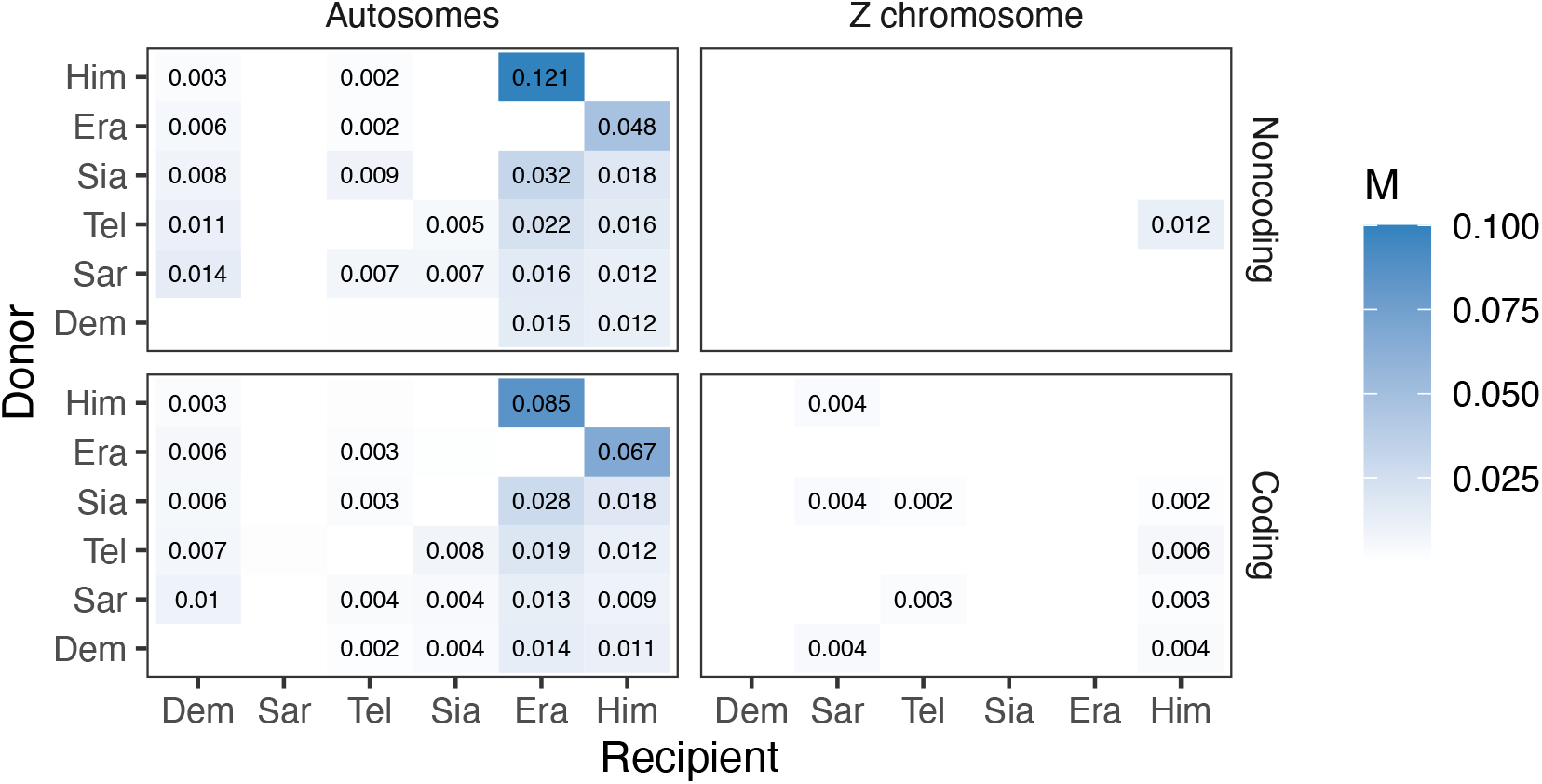
Maximum-likelihood estimates of the migration rates (*M* = *Nm*) under the IM model from 3s using all 28,204 coding and 17,428 noncoding loci in the autosomes (left column) and 1,348 coding and 574 noncoding loci in the Z chromosome (right column) for each pair of species. The donor and recipient species of gene flow are given in the y- and x-axis, respectively. *H. melpomene* was used as an outgroup. Only significant migration rate estimates (by likelihood ratio test at *p* < 0.01) larger than 0.001 are shown. See **Figure S2** and **Table S8** for estimates of other parameters.

Given tree ii, the Z chromosome (in particular the noncoding loci) was almost devoid of gene flow, in sharp contrast with the autosomes (**Fig. 3, Table S8**). For the autosomes, the highest rates of migration according to 3s were between the two sister species *H. erato* and *H. himera*, with gene flow occurring in both directions (at the rate of 0.085–0.121 migrants per generation from *H. himera* to *H. erato* and 0.048–0.067 in the opposite direction) (**Fig. 3**). There was gene flow into *H. erato, H. himera* and *H. demeter* from every other species, and from *H. sara* to all other species. These results were largely consistent among the individual autosomes (**Fig. S1**), across pairs of species (**Fig. S2**).

The pattern of gene flow inferred using 3S may reflect complex introgression in this group of species as well as the difficulty of using pairwise migration rates to reconstruct the full migration history for all species. If gene flow involved ancestral branches on the tree for all six species, we would expect the LRT to detect it in multiple pairs of species, although with distorted estimates of times and rates of migration. One scenario is extensive introgression involving common ancestors of the *sara* group and the *erato* group, which should show up as initial gene flow after species divergence that ceased after a certain time.

We used the results from the previous two analyses, blockwise BPP and 3S, to formulate a plausible history of species divergences and cross-species introgression for the *erato-sara* clade. The Z chromosome tree from the BPP analysis was used as the backbone, onto which 3s-supported introgression events were added to reconcile other species trees from the blockwise BPP analysis (**Fig. 4a**).

**Figure 4.**
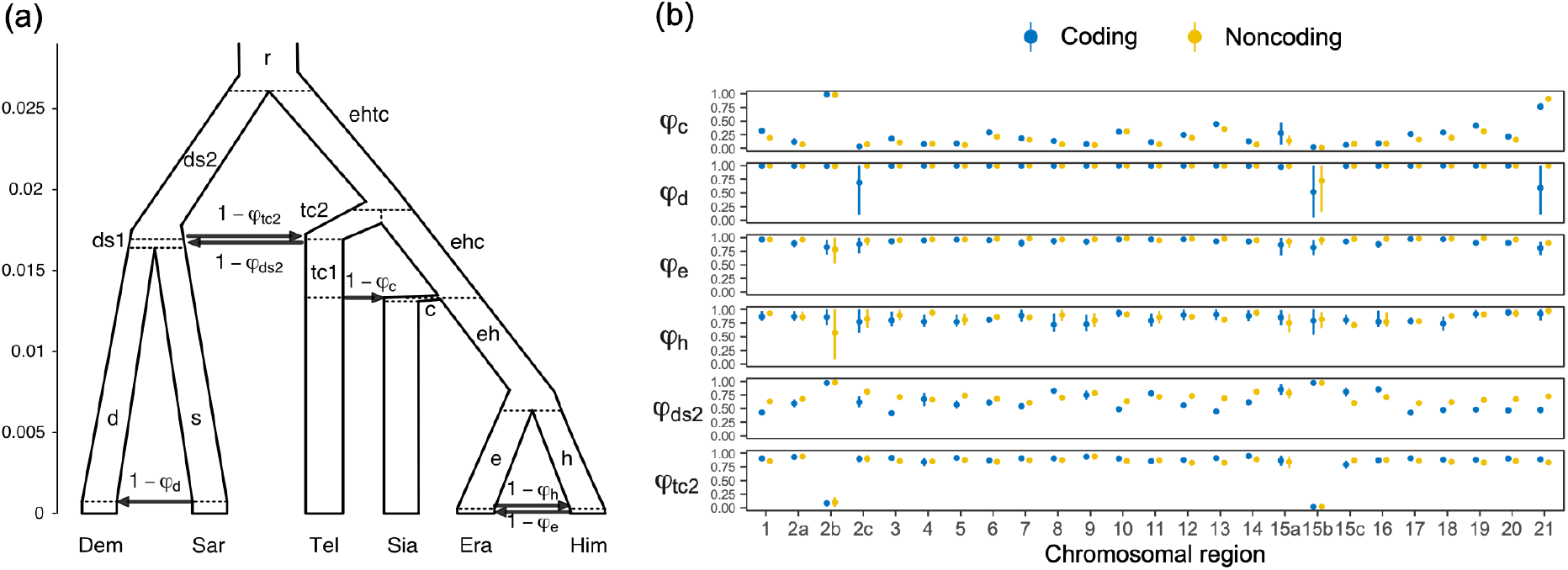
(**a**) The introgression (MSci) model, proposed based on the BPP species tree estimation under MSC and 3S analysis under the IM model, involving two unidirectional introgression events (s → d with introgression probability 1 – *φ*_d_, and tc1 → c with probability 1 – *φ*_c_) and two bidirectional introgression (BDI) events (e ↔ h with probabilities 1 – *φ*_e_ and 1 – *φ*_h_, and ds2 ↔ tc2 with probabilities 1 – *φ*_ds2_ and 1 – *φ*_tc2_). The model involves 34 parameters: 5 species divergence times and 4 introgression times (*τ*s), 19 population sizes (*θ*s), and 6 introgression probabilities. To identify parameters (see **Fig. S3**), internal nodes are labelled with lowercase letters, e.g., e is the parent node of Era, eh is the parent node of e and h, etc. Each branch corresponds to a population and is referred to using the label for its daughter node, e.g., branch e-Era is labelled branch Era and has population size *θ*_Era_, and branch eh-e is labelled branch e and has *θ*_e_. Branch lengths in the tree represent posterior means of divergence/introgression times in the BPP analysis of the 6,030 noncoding loci in chromosome 1. Estimates for other chromosomes or for coding loci are presented in **Figure 5**. (**b**) Posterior means (dots) and 95% HPD intervals (bars) of the six introgression probabilities (1 – *φ*) under the MSci model of (**a**) obtained from BPP analysis of the 25 chromosomal regions (see **Table S4** for the number of loci). Estimates for other parameters are in **Figure S3**. Inversion regions 2b and 15b on chromosomes 2 and 15 were analyzed separately, and there was an alternative set of posterior estimates resulting from within-model unidentifiability; see **Figure 6**.

### Construction of a full history of species divergences and cross-species introgression

We then used the MSci model or species tree with introgressions of **Figure 4a** to estimate the introgression probabilities (1 – *φ*) as well as species divergence/introgression times (*τ*) and population size parameters (*θ*) for each chromosomal region. (Note: in the version of BPP used here, 1 – *φ* is the introgression probability for the horizontal branch representing introgression, while more recent versions from 4.4.0 onwards use *φ* instead). All coding loci are analyzed as one dataset for each chromosomal region, as are all noncoding loci. These correspond to the A00 analysis under MSci in BPP (Flouri et al. 2020).

Estimates of introgression probability from *H. telesiphe* into *H. hecalesia* were consistently high (1 – *φ*_c_> 0.5, ∼0.8 on average) across the genome, except for the 2b inversion region and Z chromosome (**Figs. 4b, 5**). The time estimates suggest that this introgression occurred almost immediately after *H. hecalesia* split from the common ancestor of *H. erato* and *H. himera* (**Fig. 5**), supporting the hypothesis that *H. hecalesia* is a hybrid species generated during a single catastrophic event. Even though our model assumed different times for species divergence (*τ*_ehc_) and for introgression (*τ*_c_), with *τ*_ehc_> *τ*_c_(equivalent to model B in Flouri et al. (2020)), posterior estimates strongly suggest that those two times actually coincided, with *τ*_ehc_≈ *τ*_c_(equivalent to model C in Flouri et al. (2020)). This pattern was consistent with our estimates of the species trees from blockwise BPP analysis (**Fig. 2**) where the autosomes, except for the 2b region, were dominated by tree i as a result of *H. telesiphe* → *H. hecalesia* introgression on tree ii.

**Figure 5.**
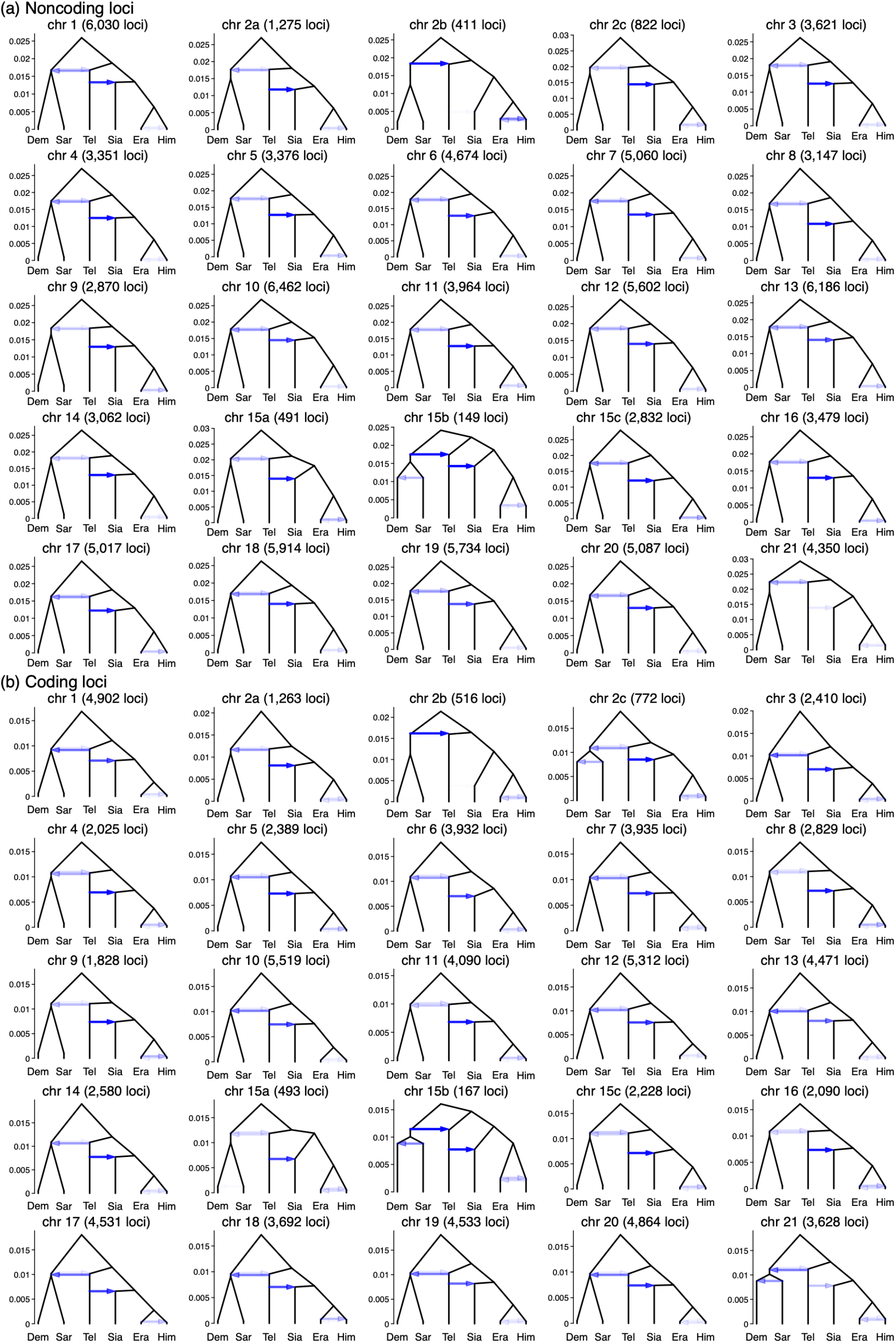
Estimated introgression history in each chromosomal region obtained from bpp analysis under the MSci model (**Fig. 4a**) using (**a**) noncoding and (**b**) coding loci. Intensity of the horizontal edges represents the posterior means of the six introgression probabilities (see **Fig. 4b**), while the y-axis represents the nine divergence/introgression times in the expected number of mutations per site. A full list of posterior estimates and 95% HPD intervals, including nineteen population size parameters, is in **Table S9**.

For the more ancient introgression between *H. telesiphe* and the common ancestor of *H. demeter* and *H. sara*, the estimated introgression probability from *H. telesiphe* (1 – *φ*_ds2_) was substantial across the entire genome, ∼0.3–0.4 on average, with 1 – *φ*_tc2_ ≈ 0.1 in the reverse direction (**Fig. 4b, Table S9**). This suggests genome-wide flow between the *erato* clade and the *sara* clade prior to the *H. telesiphe* → *H. hecalesia* introgression/hybridization. The estimates of 1 – *φ*_ds2_ and 1 – *φ*_tc2_ in the two small inversion regions, 2b and 15b, were more extreme, very close to either 0 or 1, with complex identifiability issues. We discuss the introgression history of the two inversion regions below.

Other introgression events had relatively low probabilities across the genome despite evidence from the 3S analysis under the IM model. Introgression between *H. erato* and *H. himera* had probabilities of 1 – *φ*_h_ ≈ 15% for *H. erato* → *H. himera* and 1 – *φ*_e_ ≈ 5% for *H. himera* → *H. erato*, consistently throughout the genome (**Fig. 4b**). Note that the introgression probability in the MSci model is expressed as the proportion of immigrants in the receiving population at the time of hybridization, so the smaller rate in the *H. himera* → *H. erato* direction may reflect the large population size for *H. erato*. In contrast, the migration rate in the IM model is estimated to be in the range of 0.07–0.12 migrants per generation, and is larger in the *H. himera* → *H. erato* direction (**Fig. 3**). These gene flow estimates also vary considerably among chromosomes (**Figs. S1**–**S2, Table S8**).

Lastly, our BPP MSci analysis did not support the *H. sara* → *H. demeter* introgression suggested by 3s analysis. The introgression probability (1 – *φ*_d_) was either small (<1%) or had a large posterior interval in all chromosomal regions (**Fig. 4b**), neither of which provided strong support for such introgression.

Overall, estimates of species divergence/introgression times (*τ*) and population sizes (*θ*) were broadly consistent across chromosomal regions as well as between coding and noncoding loci, with sufficient numbers of loci to yield estimates with high precision (**Fig. S3, Table S9**). In particular, estimates of divergence times were highly similar among chromosomal regions and were nearly perfectly proportional between coding and noncoding loci, with the regression *τ*_C_ ≈ 0.6*τ*_NC_(*r*^*2*^ ≥ 0.99) (**Fig. S4**). The slope suggests that purifying selection on coding loci reduced the neutral mutation rate to about 0.6 × that of noncoding loci. While the noncoding and coding regions of the genome have drastically different functions, when used as markers in the BPP analyses they yield highly consistent estimates concerning the history of species divergences and cross-species introgression. This consistency indicates that the results are reliable. Similarly, the estimates of population sizes were nearly proportional between coding and noncoding loci, with the regression *τ*_C_ ≈ 0.6*τ*_NC_(*r*^*2*^ ≥ 0.9) (**Fig. S5**). Background selection may be expected to reduce within-species polymorphism (*θ*) via linked selection but not between-species divergence (*τ*) (Shi and Yang 2018). The near identical regression slopes for both *τ* and *θ* suggest a limited role of background selection in reducing neutral sequence variation.

The age of the base of the *erato-sara* clade (*τ*_r_) was estimated to be about 0.027 substitutions per site for noncoding loci on average (**Fig. S3, Table S9**). If we use the mutation rate of 2.9 × 10^−9^ neutral mutations per site per generation and 4 generations per year (Keightley et al. 2015), this translates to about 2.3 million years (Ma) of divergence.

Population size parameters (*θ*) for populations corresponding to short branches on the species tree (**Fig. 4a**) were difficult to estimate reliably, with considerable variation among chromosomes and large posterior intervals (**Fig. S3, Table S9**), reflecting both low information content and the impact of the heavy-tailed inverse gamma prior. Most populations had *θ* in the order of 0.01 while a few had more extreme estimates. *H. erato* (*θ*_Era_) had small values, in the order of 0.0001–0.001, for most chromosomes, except in the 2b inversion where the *θ*_Era_ estimates were unreliable due to lack of data. In the current implementation of the MSci model, the same species before and after an introgression event are considered distinct species and are assigned independent *θ* parameters. It appears more sensible to assign the same *θ*, in particular when the introgression event is inferred to be nonexistent (as in the case of the *H. sara* → *H. demeter* introgression) and/or to be extremely recent. Thus we believe the estimates of *θ* for the parent populations prior to introgression events are more appropriate estimates for the four modern species: *H. demeter* (*θ*_d_ ≈ 0.01, better than *θ*_Dem_ ≈ 0.002-0.004), *H. sara* (*θ*_s_ ≈ 0.01, better than *θ*_Sar_ ≈ 0.06-0.13), *H. erato* (*θ*_e_ ≈ 0.06-0.30, better than *θ*_Era_ ≈ 0.0001-0.0166) and *H. himera* (*θ*_h_ ≈ 0.005-0.017, better than *θ*_Him_ ≈ 0.0009-0.0022) all for noncoding autosomal loci (**Table S9**). The genome sequences of *H. erato* and *H. himera* were obtained from partially inbred individuals to aid assembly, which likely explains their apparent low effective population sizes and large variation among chromosomes. The *H. erato* genome, in particular, has long stretches of homozygosity on some chromosomes due to inbreeding (**Fig. S6**) corresponding approximately to per-chromosome estimates of *θ*_Era_.

## Discussion

### Major features of the species tree and introgression: comparison with previous studies

In this study, we processed genomic sequence data from the *erato-sara* clade of *Heliconius* from Edelman et al. (2019) and compiled unphased diploid sequence alignments. We assessed the probable genotype calling error rate based on our high coverage reads and used a high coverage filter to exclude almost all genotype calling errors. We then used the data to infer the history of speciation and introgression under the MSci model in a full-likelihood framework, accounting for unphased diploid genomic data probabilistically without collapsing heterozygous sites. This coalescent-based full-likelihood method is an improvement over summary methods or sliding-window/concatenation analysis, which make less efficient use of information from genomic sequence data.

In general, our analysis confirms several major conclusions from previous analyses but reveals improved details of the evolutionary history for the clade. As in Edelman et al. (2019), we find that the Z chromosome supports tree ii, while autosomes favour tree i. We weigh the evidence and find that the Z chromosomal tree is almost certainly the true species tree (see ‘Pairwise gene flow rates’), even though large-scale introgression has distorted genealogical histories of autosomes so that the most common gene tree in the genome has a different topology (mostly tree i). Such major genealogical conflicts between the sex chromosome and the autosomes were inferred also in other species groups including *Anopheles gambiae* (Fontaine et al. 2015; Thawornwattana et al. 2018), felids (Li et al. 2019) and small finches (Stryjewski and Sorenson 2017). In the *Anopheles gambiae* group of African mosquitoes, the X chromosomal tree was similarly inferred to be the species tree while the autosomal tree reflects rampant gene flow (Fontaine et al. 2015; Thawornwattana et al. 2018). However, sliding-window analysis of genomic data from the *Anopheles gambiae* group (Fontaine et al. 2015) led to an incorrect species tree; coalescent-based full-likelihood methods produced a tree more consistent with chromosomal inversion data and with established patterns of cross-species hybridization and gene flow (Thawornwattana et al. 2018). In the *Heliconius* dataset, in contrast, concatenated windows and MSci analyses yield similar species trees. Concatenation may produce anomalous gene trees (Roch and Steel 2015) so that the estimated gene trees from large sliding windows may differ from the species phylogeny, but this was not the case for the *Heliconius* data. In the *Anopheles gambiae* group, all species arose very rapidly, creating very short internodes at the base of the species tree, while the six *Heliconius* species studied here are well separated with longer internal branches (**Fig. 4a**). Thus, the sliding-window gene tree approach was more successful in *Heliconius* than in the *Anopheles gambiae* group.

We also tested the impact of phasing errors, by generating datasets with heterozygote phase resolved at random for chromosome 1, to mimic the haploid consensus sequences. From the blockwise BPP MSC analysis (30 noncoding and 25 coding 200-locus blocks), we found that phasing errors had little impact on species tree estimation, with random phasing producing identical or highly similar posterior probabilities and the same MAP species tree. However, phasing errors had greater impact on estimation of parameters under the MSci model of **Figure 4a**: the divergence times and modern population sizes tend to be overestimated (**Fig. S7**, using all 4,902 coding loci and 6,030 noncoding loci from chromosome 1). These results are consistent with simulations, where species tree estimation was more robust and parameter estimation was more sensitive to phasing errors (Huang et al. 2021).

Our improved data processing and the use of full-likelihood approach also made it possible to answer additional questions. For example, *D* statistics and internal branch length tests (Edelman et al. 2019) can be used to test for the presence of gene flow, but cannot identify gene flow between sister species, infer the direction of gene flow or estimate its magnitude between non-sister species. In contrast, migration rates and introgression probabilities in the IM and MSci models, which measure the magnitude of gene flow, can be estimated from genomic data using full likelihood methods (Jiao et al. 2021). The proportions of gene trees in sliding-window/concatenation analyses are sensitive to the window size and are not directly comparable with introgression probabilities or even with proportions of species trees in the blockwise BPP MSC analysis.

For example, the frequencies of estimated gene trees i and ii in Edelman et al. (2019) are 24.3% and 25.8% among 10kb sliding windows (their Fig. S78), but are 35.5% and 34.8% among the 50kb windows (mapped to the *H. erato* reference, their Fig. 2B), and 36.4% and 45.4% among the 50 kb windows (mapped to the more distant *H. melpomene* reference, their Fig. S77). The proportion of windows with the dominant gene tree will increase when the window becomes larger due to the averaging effect of concatenation. In our blockwise BPP MSC analysis, local variation in the rate of gene flow among species (affected by linkage to selected loci and local recombination rate) is expected to be the most important factor leading to fluctuations among blocks, as gene-tree conflicts due to ancestral polymorphism are filtered out by the MSC model. In the BPP MSC analysis, the frequencies of (species) trees i and ii are 54.8% and 42.8% among the 100-loci noncoding blocks, 44.2% and 48.3% among the 100-loci coding blocks, 58.9% and 40.2% among the 200-loci noncoding blocks, and 46.9% and 49.5% among the 200-loci coding blocks. These are reasonably consistent between coding and noncoding loci and seem less sensitive to the number of loci in each block than is the sliding-window analysis to the window size.

There was a greater representation of the presumed introgression tree i in noncoding loci (55-59%) than in coding loci (44-47%) compared with tree ii, and a moderate excess of *H. telesiphe* → *H. hecalesia* introgression (1 – *φ*_c_) in noncoding versus coding loci on each chromosome (**Fig. 4b**). This may reflect greater constraint on introgression for coding sequences as found in other systems, for instance for Neanderthal ancestry in modern humans (Sankararaman et al. 2014).

Previous estimates of migration rates and timings in *Heliconius* under models of gene flow such as the IM model have been limited to a few loci (Bull et al. 2006; Kronforst et al. 2006; Kronforst 2008; Salazar et al. 2008; Pardo-Diaz et al. 2012) or a few species (Kronforst et al. 2013; Van Belleghem et al. 2021), most of which have focused on the *melpomene* clade. Using joint site-frequency spectrum and a secondary contact model, Van Belleghem et al. (2021) estimated migration rates (*M* = *Nm*) to be around 0.5–0.6 migrants per generation from two races of *H. erato* (*H. erato favorinus* and *H. erato emma*) to *H. himera*, and 0.07–0.13 in the opposite direction. This asymmetric gene flow accords with our MSci results, in which the probability of flow is 15% from *H. erato* to *H. himera*, and 5% in the opposite direction (**Fig. 4b, Table S9**). Our estimates from the 3S analysis of the genomic data are 0.05–0.07 migrants per generation from *H. erato* to *H. himera*, and 0.09–0.12 in the opposite direction. The differences may be due to different data and methods used. No estimates of other introgression probabilities in **Fig. 4** have previously been reported.

### Limitations of our analyses

Implementation of the MSci model in BPP (Flouri et al. 2020) provides a powerful tool for estimating the timing and magnitude of introgression on a species phylogeny from genome-scale data while accounting for deep coalescence and genealogical heterogeneity across the genome. The implementation also retains heterozygous sites in the genomic data and averages over all possible heterozygote phase resolutions (Gronau et al. 2011), avoiding systematic errors caused by the use of haploid consensus sequences (Huang et al. 2021). Other likelihood-based methods incorporating cross-species gene flow are under active development (Hey et al. 2018; Wen and Nakhleh 2018; Zhang et al. 2018; Jones 2019), but they are computationally demanding and currently impractical for data of >100 loci. We have also tested for the effects of sequencing error on genotype calls; erroneous genotype calls may also potentially obscure the genealogical signal in the data. We find that genotype call error in our data should be very low due to high read coverage (>20x after processing).

Here we discuss several limitations of our approach. First, the introgression model we formulated (**Fig. 4a**) relies on two main sources of information: variation in the species tree estimated from blocks of loci across the genome and pairwise migration rate estimates from 3S. Integrating such information into a single MSci model is non-trivial. Ideally, we would be able to infer the species phylogeny and introgression events simultaneously with associated parameters such as introgression and divergence times, but this has yet to be implemented.

Second, even with a fixed MSci model (**Fig. 4a**), inference of demographic parameters is computationally challenging. Some parameters were difficult to estimate reliably, such as population sizes associated with short branches in the species tree (e.g., *θ*_Sar_ and *θ*_ds1_; see **Fig. S3**). Contributing factors include unidentifiability of the model, multimodal posterior distribution, poor MCMC mixing, and potential model misspecification. The unidentifiability arises because of the symmetry in the likelihood induced by bidirectional introgression (BDI, such as the two BDI events in the model of **Figure 4a**) (Flouri et al. 2020). The BDI between nodes ds2-tc2 (**Fig. 4a**) is between non-sister species and creates a pair of unidentifiable models (**Fig. S8**). The BDI at nodes e-h (**Fig. 4a**) is between sister-species and leads to a pair of unidentifiable likelihood peaks within the same model (not shown). We prefer species tree *S*_1_ of **Figure S8** because the introgression probabilities are 1 – *φ*_ds2_< ½ and 1 – *φ*_tc2_ ≈ 0.1, while the alternative model *S*_2_ has introgression probabilities *φ*_ds2_> ½ and *φ*_tc2_ ≈ 0.9, implying mutual replacements of the species involved; this does not seem plausible biologically. Similarly for the BDI event at e-h, we focus on the peak with small introgression probabilities (with 1 – *φ*_e_< ½ and 1 – *φ*_h_< ½) (**Fig. 4b**). In other words, we resolve the unidentifiability caused by BDI events by applying simple constraints such as 1 – *φ* < ½, as recommended by Flouri et al. (2020). An example of MCMC outputs and processing is provided in **Figure S9**. Furthermore, unidentifiability can occur when the introgression probability is estimated to be 0 or 1. This is the case for the 2b and 15b inversions; see our discussions of the different inversion introgression scenarios below.

Third, we avoid explicit modeling of recombination by analyzing short genomic loci (∼200 bp on average) that are far apart (at least 2 kb). This assumes that sites within each locus have zero recombination, whereas different loci are free to recombine. Sites are expected to be approximately independent at physical distances of ∼10 kb apart, at least based on linkage disequilibrium decay estimated from the *H. melpomene* group species (Dasmahapatra et al. 2012, their Fig. S16.2.1). Thus, some correlation of gene genealogies between consecutive loci may be expected. The unrealistic assumption of free recombination between loci is expected to have a slight effect of exaggerating information content within each block of loci, leading potentially to artificially narrow credibility intervals for parameter estimates. The assumption of no recombination among sites of the same locus may be of more concern since recombination causes different parts of the sequence to have different histories while the model assumes one history. However, the loci used in this study are so short that the assumption of no recombination within a locus is expected to hold approximately. Previously, Lanier and Knowles (2012) found that species tree estimation under MSC is robust to realistic levels of recombination. The impact of recombination on estimates of the migration rate or introgression probability is not well understood, in particular when recombination rate varies across the genome.

Fourth, the migration (IM) models in 3S and the MSci model in BPP are necessarily simplified. Currently we lack a good understanding of the behaviour of our inference methods when the model of gene flow is misspecified, for example, when the MSci model is fitted to data generated under the IM model or vice versa (Jiao et al. 2020, Fig. 7). One may expect the magnitude of gene flow to vary across the genome and over time. We are currently investigating the performance of estimation methods such as 3S and BPP under such complex scenarios of gene flow, in order to develop methods that account for such variation.

### Introgression history of the two inversion regions

Our analyses consistently suggest that two inversion regions, 2b and 15b, have genetic histories distinct from the rest of the genome (**Figs. 4b, 5**). All introgression probabilities for those two regions were close to either 0 or 1, except for estimates which involve large uncertainties due to lack of data. This suggests that loci within each of the two inversion regions were inherited almost without recombination as a single locus. This is likely if an inverted region introgresses and becomes fixed in a population that previously lacked the inversion (or vice-versa), and if, during the period of polymorphism, recombination in the inverted region was strongly suppressed – a known biological effect of inversions. Although the MSC framework explicitly models ancestral polymorphism, inversion polymorphism is not accounted for since it creates substructure within the ancestral population while the model assumes random mating within species and free recombination among loci. Nonetheless, introgression probabilities of either 0 or 1 are sensible estimates. When introgression probabilities are near 0 or 1, some parameters can become unidentifiable, and different parameter values corresponding to different introgression histories can explain the data equally well. For each of the two inversion regions, we found distinct peaks in the posterior distribution which correspond to different historical scenarios with opposite introgression directions between (*H. sara, H. demeter*) and *H. telesiphe* (**Fig. 6**). Both scenarios are consistent with the known inversion orientations in modern species. Here, we discuss evidence in favour of each introgression scenario, although it is difficult to rule out alternatives.

**Figure 6.**
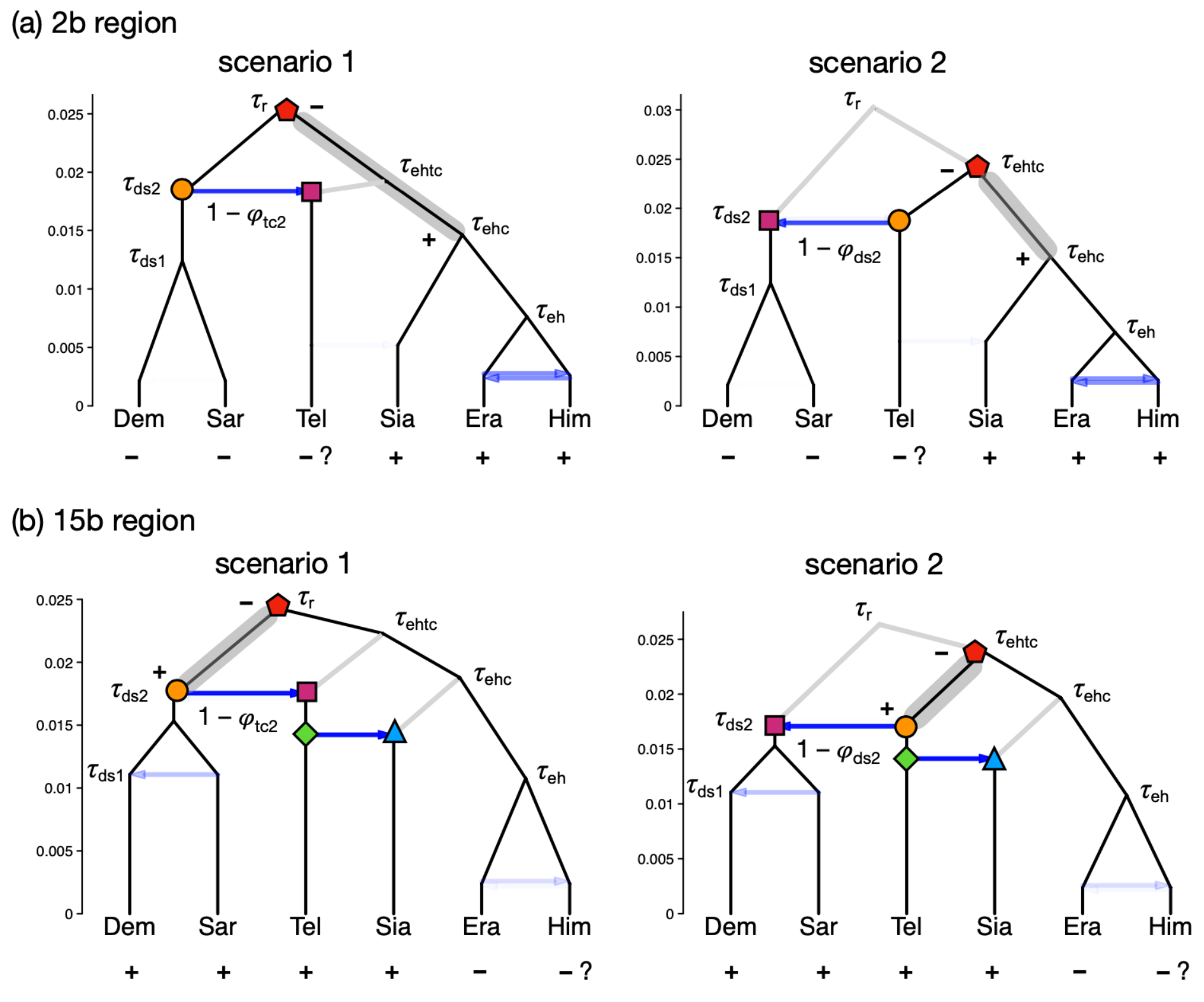
Within-model unidentifiability and possible introgression histories for inversion regions (**a**) 2b and (**b**) 15b, showing estimated divergence times from BPP analysis under the MSci model (**Fig. 4a**). For each region, there are two competing scenarios of the introgression history that fit the data equally well, resulting in unidentifiability of certain model parameters. Plus (+) represents the derived inverted orientation while minus (–) is the ancestral non-inverted orientation. Species with uncertain inversion orientation are marked with ‘?’. Node symbols and colours indicate matching nodes and times between the two scenarios. The estimates were based on posterior means from noncoding loci. Grey band indicates an ancestral population within which the derived inverted orientation (+) may have arisen from the non-inverted orientation (–). Note that parameters associated with nodes reached by very few sequences (indicated as grey branches) were expected to be poorly estimated, with a wide posterior interval. For example, in scenario 1 for 2b, *τ*_ehtc_, *θ*_ehtc_ and *θ*_ehc_ were poorly estimated (**Fig. S3, Table S9**).

In the chromosome 2b region (1.95 Mb, with 411 noncoding loci and 516 coding loci), introgression between the common ancestor of the *sara* clade and *H. telesiphe* was inferred to be unidirectional with 1 – *φ*_tc2_ ≈ 0.9 and 1 – *φ*_ds2_ ≈ 0 in scenario 1 (**Fig. 6a**), while in scenario 2, introgression was mostly unidirectional but in the opposite direction (1 – *φ*_tc2_ ≈ 0, 1 – *φ*_ds2_ ≈ 0.8). Both scenarios are consistent with tree iii (**Fig. 2**), where *H. telesiphe* clusters with the *sara* clade and the *H. telesiphe* → *H. hecalesia* introgression was absent (1 – *φ*_c_ ≈ 0). The two scenarios make similar predictions about the gene-tree histories and the genetic data (for example, both scenarios predict the origin of the inversion in the branch ehc), but the effective root of the species tree for the 2b region is at node r in scenario 1 and at node ehtc in scenario 2.

To determine which scenario is more consistent with other autosomal regions of the genome, we compared the root age of the inversion region with *τ*_r_ and *τ*_ehtc_ on other chromosomes. Divergence times (*τ*) for the 2b region are highly similar to those in other regions of the genome, with regression slopes ∼1.01–1.08 for the noncoding loci (**Fig. S10**), suggesting that the 2b region has a neutral mutation rate similar to that of other genomic regions. Scenario 1 is supported if the root age (*τ*_r_ in scenario 1, *τ*_ehtc_ in scenario 2) for inversion 2b is close to *τ*_r_ in other regions, while scenario 2 is supported if it is closer to *τ*_ehtc_ from the rest of the genome. For noncoding loci, the root age of 2b (posterior mean 0.026) was closer to the root age of the rest of the genome expected under scenario 1 (*τ*_r_= 0.026–0.029) than that for scenario 2 (*τ*_ehtc_= 0.018–0.023, **Fig. S3, Table S9**). Although we do not know whether *H. telesiphe* has the inversion, its ancestor likely had the inversion, but introgression from the ancestor of *H. demeter* and *H. sara* (which lack the inversion) under scenario 1 almost certainly caused replacement without recombination. We therefore predict that *H. telesiphe* does not today carry the inversion, but that its ancestor did. In contrast, scenario 2 predicts that *H. telesiphe* always lacked the inversion, but it would be unclear why the non-inverted region from *H. telesiphe* replaced the original non-inverted sequence in the *H. demeter/H. sara* ancestor with an almost complete lack of recombination. Thus, this secondary evidence also supports scenario 1 (**Fig. 6a**).

An alternative to an introgression hypothesis requires ancestral polymorphism of the 2b inversion, where *H. sara, H. demeter* and *H. telesiphe* retained the ancestral non-inverted orientation through incomplete lineage sorting after divergence of the *erato* and *sara* clades (Edelman et al. 2019). Compared with the introgression scenario, ancestral inversion polymorphism predicts that the divergence time *τ*_tc2_ for the common ancestor of *H. telesiphe, H. sara* and *H. demeter* must pre-date or overlap the divergence time of the entire clade found for the rest of the genome. Our results did not support this hypothesis. Estimated *τ*_tc2_ in the 2b region (for noncoding regions, 0.018) was always much closer to the introgression time in the rest of the genome (*τ*_tc2_= 0.016–0.022) than to the root in the rest of the genome (*τ*_r_= 0.026–0.029) (**Fig. S3, Table S9**), suggesting that the introgression hypothesis is more likely than the inversion polymorphism hypothesis in the ancestral population prior to the divergence of the entire *erato-sara* clade.

Based on pairwise sequence distances (*D*_*XY*_), Edelman et al. (2019) argued that ancestral inversion polymorphism could explain the data as well as or better than (*H. sara, H. demeter*) → *H. telesiphe* introgression for inversion 2b (their Fig. S95). The introgression scenario was rejected because the *H. sara*–*H. telesiphe* divergence of 2b was not much lower than the genome-wide average. However, this conclusion assumes that that all other chromosomes were free of such introgression. The prevalence of BDI between (*H. sara, H. demeter*) and *H. telesiphe* across the genome, as suggested by our results (**Figs. 4b, 5**), could explain the patterns of *D*_*XY*_ in Edelman et al. (2019). The possibility of the opposite introgression with *H. telesiphe* as the source species (scenario 2 in **Fig. 6a**) was also not considered by Edelman et al. (2019).

For the chromosome 15b inversion region (580 kb, with only 149 noncoding loci and 167 coding loci), two sets of estimates were always obtained from independent MCMC runs, each occurring about 50% of the time. This inversion contains the *cortex* gene known to be involved in mimicry in the *H. melpomene*/silvaniform group as well as the *H. erato*/*sara* group of *Heliconius* (Nadeau et al. 2016), although the precise role of the inversion in mimicry is unknown for the species studied here (Edelman et al. 2019). As for the 2b region, a pair of scenarios correspond to alternative histories differing in direction of introgression between the common ancestor of the *sara* clade and *H. telesiphe* (**Fig. 6b**). Again, both scenarios are consistent with the pattern of shared derived orientation of the inversion region among *H. telesiphe, H. hecalesia, H. demeter* and *H. sara*, as represented by tree iv (**Fig. 2**). In both scenarios, the inversion originated along the branch between the root and the introgression between the ancestral populations of *H. telesiphe* and *H. demeter/H. sara* (**Fig. 6b**). As before, we used the root age (*τ*_r_ under scenario 1 and *τ*_ehtc_ under scenario 2) to distinguish between these two possibilities. Note that the divergence time estimates in the 15b region are also highly comparable to those in other genomic regions, with regression slopes ∼1.00– 1.07 for the noncoding loci (**Fig. S11**). The comparison suggests that introgression from the common ancestor of *H. sara* and *H. demeter* into the ancestor of *H. telesiphe* was more compatible with the other regions of the genome than the opposite direction (**Fig. S3, Table S9**), even though the opposite direction of introgression was marginally more prevalent across the rest of the genome (**Fig. 5**). Our result for 15b is concordant with the findings of Edelman et al. (2019), who used internal branch lengths and *D*_*XY*_ to suggest introgression as a likely scenario, although they did not consider the alternative scenario 2 of introgression (**Fig. 6b**).

Our results provide evidence that introgression according to scenario 1 (**Fig. 6**) between (*H. sara, H. demeter*) and *H. telesiphe* rather than ancestral polymorphism is a more likely explanation for the history of both 2b and 15b inversion regions, and that the introgression was almost wholly unidirectional. Additional genomes with inversion status from diverse species from this *erato-sara* clade and better breakpoint characterization will be useful to test these conclusions about the alternative scenarios.

### Evidence of introgression from natural populations

Our analyses support three introgression events (**Fig. 4**): (1) between *H. telesiphe* and the common ancestor of the *sara* clade, (2) from *H. telesiphe* into *H. hecalesia*, and (3) between *H. erato* and *H. himera*. The first two are consistent with previous genomic studies (Edelman et al. 2019; Kozak et al. 2021). Ancient introgression such as (1) will be difficult to confirm using evidence from natural populations today. The recent introgression (3) is well-documented in natural populations and in mating experiments (Jiggins et al. 1997; Mcmillan et al. 1997; Mallet et al. 2007).

However, the six species included in this study constitute but a fraction of the species and geographic races in the *erato-sara* clade, so caution should be exercised in interpretation of inferred introgression. In particular, signals of introgression may be indirect, involving related species or subspecies unsampled in the data.

For example, the *H. telesiphe* → *H. hecalesia* introgression inferred from our data may not represent direct gene flow between the two species, which do not currently overlap geographically (Rosser et al. 2012). *H. hecalesia* does overlap with *H. clysonymus* and its sister species *H. hortense* in West Ecuador, the Colombian Andes, and Central America (Rosser et al. 2012), and there are documented natural hybrids between *H. hecalesia* and both *H. clysonymus* and *H. hortense* (Mallet et al. 2007). Furthermore, *H. telesiphe* forms a well-supported clade with *H. clysonymus* and *H. hortense*, nested within the *erato* clade (Kozak et al. 2015; Massardo et al. 2020; Kozak et al. 2021). Therefore, it is likely that introgressed loci in *H. hecalesia* actually came from *H. clysonymus, H. hortense*, or their common ancestor. Introgression from *H. clysonymus* or *H. hortense* is also supported by phylogenetic network analysis and *D* statistics in Kozak et al. (2021).

Another case is the possible indirect gene flow between *H. erato* and *H. himera. H. himera* is considered an incipient species within *H. erato* (*sensu lato*). It is restricted to middle elevations (800–2,000 metres) of the Andes in South America; in contrast, the subspecies of *H. erato* characterized here, *H. erato demophoon*, is found in Central America (Rosser et al. 2012). However, *H. himera* is parapatric with several subspecies of *H. erato* (such as *cyrbia, favorinus, lativitta* and *emma*) (Mallet 1993; Rosser et al. 2012), with narrow contact zones where natural hybrids can be found at frequencies of 5–10% (Jiggins et al. 1996; Jiggins et al. 1997; Mallet et al. 1998). The recent introgression signal we detect here almost certainly resulted from these adjacent subspecies of *H. erato* rather than from *H. erato demophoon*.

### Conclusion

The coalescent-based full-likelihood approach employed here was able not only to detect gene flow among species, but also to estimate its direction, timing and magnitude using genomic data. We have demonstrated that a much fuller understanding of the demographic and introgression history of a well-studied group of species can be gained by such an approach than with methods more usually employed hitherto. In the *Heliconius erato-sara* clade of species, we found robust evidence of introgression across the genome involving distantly related species deep in the phylogeny as well as between sister species in shallower parts of the tree. We confirm ancestral gene flow between the *sara* clade and an ancestor of *H. telesiphe*, infer a likely hybrid speciation origin of *H. hecalesia*, as well as gene flow between the sister species *H. erato* and *H. himera*. We clarify how introgression among ancestral species can explain the history of two chromosomal inversions deep in the phylogeny of the *erato-sara* group. Importantly, we estimate key population parameters such as species divergence times and population sizes for modern and ancestral species of this *Heliconius* clade, providing an opportunity to understand the speciation and introgression history of this group in fine detail.

## Supplementary Material

Supplementary materials are available in Zenodo at https://doi.org/10.5281/zenodo.5639614. The multilocus sequence datasets for BPP and 3s analyses are available in Dryad at https://doi.org/10.5061/dryad.dfn2z3526. The C program for calculating the genotyping error rate given the base-calling error and read depth is at https://github.com/abacus-gene/genotypecall. All code used to filter the data are available at https://github.com/fernandoseixas/EratoCladePhylogeny.

## Acknowledgements

We thank Nathaniel Edelman, Paul Frandsen, Mia Miyagi, and Bernardo Clavijo for their inputs to the overall project. We are grateful for the efforts of two anonymous reviewers, the Associate Editor (Jacob Esselstyn) and the Editor (Bryan Carstens), who made constructive comments; these have led to significant improvements in the presentation of our results.

## References

Andermann T, Fernandes AM, Olsson U, Töpel M, Pfeil B, Oxelman B, Aleixo A, Faircloth BC, Antonelli A. 2019. Allele Phasing Greatly Improves the Phylogenetic Utility of Ultraconserved Elements. Syst Biol. 68(1):32–46.

Balaban NQ, Helaine S, Lewis K, Ackermann M, Aldridge B, Andersson DI, Brynildsen MP, Bumann D, Camilli A, Collins JJ, et al. 2019. Definitions and guidelines for research on antibiotic persistence. Nat Rev Microbiol. 17(7):441–448.

Barton NH. 2006. Evolutionary Biology: How Did the Human Species Form? Curr Biol. 16(16):R647–R650.

Bates HW. 1862. Contributions to an Insect Fauna of the Amazon Valley. Lepidoptera: Heliconinæ. Trans Linn Soc London. 23:495–566.

Van Belleghem SM, Cole JM, Montejo-Kovacevich G, Bacquet CN, McMillan WO, Papa R, Counterman BA. 2021. Selection and isolation define a heterogeneous divergence landscape between hybridizing Heliconius butterflies. Evolution (N Y). 75(9):2251–2268.

Van Belleghem SM, Rastas P, Papanicolaou A, Martin SH, Arias CF, Supple MA, Hanly JJ, Mallet J, Lewis JJ, Hines HM, et al. 2017. Complex modular architecture around a simple toolkit of wing pattern genes. Nat Ecol Evol. 1(3):52.

Beltrán M, Jiggins CD, Brower AVZ, Bermingham E, Mallet J. 2007. Do pollen feeding, pupal-mating and larval gregariousness have a single origin in Heliconius butterflies? Inferences from multilocus DNA sequence data. Biol J Linn Soc. 92(2):221–239.

Beltrán M, Jiggins CD, Bull V, Linares M, Mallet J, McMillan WO, Bermingham E. 2002. Phylogenetic Discordance at the Species Boundary: Comparative Gene Genealogies Among Rapidly Radiating Heliconius Butterflies. Mol Biol Evol. 19(12):2176–2190.

Brower AVZ. 1994. Phylogeny of Heliconius Butterflies Inferred from Mitochondrial DNA Sequences (Lepidoptera: Nymphalidae). Mol Phylogenet Evol. 3(2):159–174.

Brower AVZ, Egan MG. 1997. Cladistic analysis of Heliconius butterflies and relatives (Nymphalidae: Heliconiiti): a revised phylogenetic position for Eueides based on sequences from mtDNA and a nuclear gene. Proc R Soc London Ser B Biol Sci. 264(1384):969–977.

Bull V, Beltrán M, Jiggins CD, McMillan WO, Bermingham E, Mallet J. 2006. Polyphyly and gene flow between non-sibling Heliconius species. BMC Biol. 4(1):11.

Dalquen DA, Zhu T, Yang Z. 2017. Maximum likelihood implementation of an isolation-with-migration model for three species. Syst Biol. 66(3):379–398.

Dasmahapatra KK, Walters JR, Briscoe AD, Davey JW, Whibley A, Nadeau NJ, Zimin A V., Salazar C, Ferguson LC, Martin SH, et al. 2012. Butterfly genome reveals promiscuous exchange of mimicry adaptations among species. Nature. 487(7405):94–98.

Davey JW, Barker SL, Rastas PM, Pinharanda A, Martin SH, Durbin R, McMillan WO, Merrill RM, Jiggins CD. 2017. No evidence for maintenance of a sympatric Heliconius species barrier by chromosomal inversions. Evol Lett. 1(3):138–154.

DePristo MA, Banks E, Poplin R, Garimella K V., Maguire JR, Hartl C, Philippakis AA, Del Angel G, Rivas MA, Hanna M, et al. 2011. A framework for variation discovery and genotyping using next-generation DNA sequencing data. Nat Genet. 43(5):491–501.

Edelman NB, Frandsen PB, Miyagi M, Clavijo B, Davey J, Dikow RB, García-Accinelli G, Van Belleghem SM, Patterson N, Neafsey DE, et al. 2019. Genomic architecture and introgression shape a butterfly radiation. Science. 366(6465):594–599.

Edelman NB, Mallet J. 2021. The prevalence and adaptive impact of introgression. Annu Rev Genet. in press.

Feder JL, Egan SP, Nosil P. 2012. The genomics of speciation-with-gene-flow. Trends Genet. 28(7):342–350.

Figueiró H V, Li G, Trindade FJ, Assis J, Pais F, Fernandes G, Santos SHD, Hughes GM, Komissarov A, Antunes A, et al. 2017. Genome-wide signatures of complex introgression and adaptive evolution in the big cats. Sci Adv. 3(7):e1700299.

Flouri T, Jiao X, Rannala B, Yang Z. 2018. Species tree inference with BPP using genomic sequences and the multispecies coalescent. Mol Biol Evol. 35(10):2585–2593.

Flouri T, Jiao X, Rannala B, Yang Z. 2020. A bayesian implementation of the multispecies coalescent model with introgression for phylogenomic analysis. Mol Biol Evol. 37(4):1211–1223.

Fontaine MC, Pease JB, Steele A, Waterhouse RM, Neafsey DE, Sharakhov I V., Jiang X, Hall AB, Catteruccia F, Kakani E, et al. 2015. Extensive introgression in a malaria vector species complex revealed by phylogenomics. Science. 347(6217):1258524.

Gronau I, Hubisz MJ, Gulko B, Danko CG, Siepel A. 2011. Bayesian inference of ancient human demography from individual genome sequences. Nat Genet. 43(10):1031–1035.

Heled J, Drummond AJ. 2010. Bayesian Inference of Species Trees from Multilocus Data. Mol Biol Evol. 27(3):570–580.

Hey J. 2010. Isolation with migration models for more than two populations. Mol Biol Evol. 27(4):905–920.

Hey J, Chung Y, Sethuraman A, Lachance J, Tishkoff S, Sousa VC, Wang Y. 2018. Phylogeny estimation by integration over isolation with migration models. Mol Biol Evol. 35(11):2805–2818.

Hey J, Nielsen R. 2004. Multilocus methods for estimating population sizes, migration rates and divergence time, with applications to the divergence of Drosophila pseudoobscura and D. persimilis. Genetics. 167(2):747–760.

Hines HM, Counterman BA, Papa R, De Moura PA, Cardoso MZ, Linares M, Mallet J, Reed RD, Jiggins CD, Kronforst MR, et al. 2011. Wing patterning gene redefines the mimetic history of Heliconius butterflies. Proc Natl Acad Sci U S A. 108(49):19666–19671.

Huang J, Bennett J, Flouri T, Leaché AD, Yang Z. 2021. Phase Resolution of Heterozygous Sites in Diploid Genomes is Important to Phylogenomic Analysis under the Multispecies Coalescent Model. Syst Biol.:2021.03.29.437575.

Jay P, Whibley A, Frézal L, Rodríguez de Cara MÁ, Nowell RW, Mallet J, Dasmahapatra KK, Joron M. 2018. Supergene Evolution Triggered by the Introgression of a Chromosomal Inversion. Curr Biol. 28(11):1839-1845.e3.

Jiao X, Flouri T, Rannala B, Yang Z. 2020. The Impact of Cross-Species Gene Flow on Species Tree Estimation. Syst Biol. 69(5):830–847.

Jiao X, Flouri T, Yang Z. 2021. Multispecies coalescent and its applications to infer species phylogenies and cross-species gene flow. Natl Sci Rev.

Jiggins CD, McMillan WO, King P, Mallet J. 1997. The maintenance of species differences across a Heliconius hybrid zone. Heredity (Edinb). 79(5):495–505.

Jiggins CD, McMillan WO, Neukirchen W, Mallet J. 1996. What can hybrid zones tell us about speciation? The case of Heliconius erato and H. himera (Lepidoptera: Nymphalidae). Biol J Linn Soc. 59(3):221–242.

Jones GR. 2019. Divergence Estimation in the Presence of Incomplete Lineage Sorting and Migration. Syst Biol. 68(1):19–31.

Keightley PD, Pinharanda A, Ness RW, Simpson F, Dasmahapatra KK, Mallet J, Davey JW, Jiggins CD. 2015. Estimation of the Spontaneous Mutation Rate in Heliconius melpomene. Mol Biol Evol. 32(1):239–243.

Kozak KM, Joron M, McMillan WO, Jiggins CD. 2021. Rampant genome-wide admixture across the <i>Heliconius<\i> radiation. Genome Biol Evol.

Kozak KM, Wahlberg N, Neild AFE, Dasmahapatra KK, Mallet J, Jiggins CD. 2015. Multilocus species trees show the recent adaptive radiation of the mimetic heliconius butterflies. Syst Biol. 64(3):505–524.

Kronforst MR. 2008. Gene flow persists millions of years after speciation in Heliconius butterflies. BMC Evol Biol. 8(1):98.

Kronforst MR, Hansen MEB, Crawford NG, Gallant JR, Zhang W, Kulathinal RJ, Kapan DD, Mullen SP. 2013. Hybridization Reveals the Evolving Genomic Architecture of Speciation. Cell Rep. 5(3):666–677.

Kronforst MR, Young LG, Blume LM, Gilbert LE. 2006. Multilocus Analyses of Admixture and Introgression Among Hybridizing Heliconius Butterflies. Evolution (N Y). 60(6):1254–1268.

Lanier HC, Knowles LL. 2012. Is recombination a problem for species-tree analyses? Syst Biol. 61(4):691–701.

Larget BR, Kotha SK, Dewey CN, Ané C. 2010. BUCKy: Gene tree/species tree reconciliation with Bayesian concordance analysis. Bioinformatics. 26(22):2910–2911.

Li G, Figueiró H V., Eizirik E, Murphy WJ, Yoder A. 2019. Recombination-Aware Phylogenomics Reveals the Structured Genomic Landscape of Hybridizing Cat Species. Mol Biol Evol. 36(10):2111–2126.

Li H. 2011. A statistical framework for SNP calling, mutation discovery, association mapping and population genetical parameter estimation from sequencing data. Bioinformatics. 27(21):2987–2993.

Li H. 2013. Aligning sequence reads, clone sequences and assembly contigs with BWA-MEM. ArXiv e-prints.

Li H, Handsaker B, Wysoker A, Fennell T, Ruan J, Homer N, Marth G, Abecasis G, Durbin R. 2009. The Sequence Alignment/Map format and SAMtools. Bioinformatics. 25(16):2078–2079.

Liu L, Yu L, Edwards S V. 2010. A maximum pseudo-likelihood approach for estimating species trees under the coalescent model. BMC Evol Biol. 10(1):302.

Malinsky M, Svardal H, Tyers AM, Miska EA, Genner MJ, Turner GF, Durbin R. 2018. Whole-genome sequences of Malawi cichlids reveal multiple radiations interconnected by gene flow. Nat Ecol Evol. 2(12):1940–1955.

Mallet J. 1993. Speciation, raciation, and color pattern evolution in Heliconius butterflies: evidence from hybrid zones. In: Hybrid zones and the evolutionary process. p. 226–260.

Mallet J, Beltrán M, Neukirchen W, Linares M. 2007. Natural hybridization in heliconiine butterflies: The species boundary as a continuum. BMC Evol Biol. 7(1):28.

Mallet J, Besansky N, Hahn MW. 2016. How reticulated are species? BioEssays. 38(2):140–149.

Mallet J, McMillan WO, Jiggins CD. 1998. Estimating the Mating Behavior of a Pair of Hybridizing Heliconius Species in the Wild. Evolution (N Y). 52(2):503.

Martin M. 2011. Cutadapt removes adapter sequences from high-throughput sequencing reads. EMBnet.journal. 17(1):10.

Martin SH, Van Belleghem SM. 2017. Exploring evolutionary relationships across the genome using topology weighting. Genetics. 206(1):429–438.

Massardo D, Vankuren NW, Nallu S, Ramos RR, Ribeiro PG, Silva-Brandão KL, Brandão MM, Lion MB, Freitas AVL, Cardoso MZ, et al. 2020. The roles of hybridization and habitat fragmentation in the evolution of Brazil’s enigmatic longwing butterflies, Heliconius nattereri and H. hermathena. BMC Biol. 18(1):84.

McKenna A, Hanna M, Banks E, Sivachenko A, Cibulskis K, Kernytsky A, Garimella K, Altshuler D, Gabriel S, Daly M, et al. 2010. The genome analysis toolkit: A MapReduce framework for analyzing next-generation DNA sequencing data. Genome Res. 20(9):1297–1303.

Mcmillan WO, Jiggins CD, Mallet J. 1997. What initiates speciation in passion-vine butterflies? Proc Natl Acad Sci U S A. 94(16):8628–8633.

Mirarab S, Reaz R, Bayzid MS, Zimmermann T S. Swenson M, Warnow T. 2014. ASTRAL: Genome-scale coalescent-based species tree estimation. Bioinformatics. 30(17):i541.-i548.

Müller F. 1879. Ituna and Thyridia; a remarkable case of mimicry in butterflies (transl. by Ralph Meldola from the original German article in Kosmos, May 1879, p. 100). Trans Entomol Soc London.:xx–xxix.

Nadeau NJ, Martin SH, Kozak KM, Salazar C, Dasmahapatra KK, Davey JW, Baxter SW, Blaxter ML, Mallet J, Jiggins CD. 2013. Genome-wide patterns of divergence and gene flow across a butterfly radiation. Mol Ecol. 22(3):814–826.

Nadeau NJ, Pardo-Diaz C, Whibley A, Supple MA, Saenko S V., Wallbank RWR, Wu GC, Maroja L, Ferguson L, Hanly JJ, et al. 2016. The gene cortex controls mimicry and crypsis in butterflies and moths. Nature. 534(7605):106–110.

Okonechnikov K, Conesa A, García-Alcalde F. 2016. Qualimap 2: Advanced multi-sample quality control for high-throughput sequencing data. Bioinformatics. 32(2):292–294.

Pardo-Diaz C, Salazar C, Baxter SW, Merot C, Figueiredo-Ready W, Joron M, McMillan WO, Jiggins CD. 2012. Adaptive introgression across species boundaries in Heliconius butterflies. PLoS Genet. 8(6):e1002752.

Patterson N, Moorjani P, Luo Y, Mallick S, Rohland N, Zhan Y, Genschoreck T, Webster T, Reich D. 2012. Ancient admixture in human history. Genetics. 192(3):1065–1093.

Payseur BA, Rieseberg LH. 2016. A genomic perspective on hybridization and speciation. Mol Ecol. 25(11):2337–2360.

Pinho C, Hey J. 2010. Divergence with gene flow: Models and data. Annu Rev Ecol Evol Syst. 41(1):215–230.

Rannala B, Yang Z. 2017. Efficient Bayesian species tree inference under the multispecies coalescent. Syst Biol. 66(5):823–842.

Reed RD, Papa R, Martin A, Hines HM, Counterman BA, Pardo-Diaz C, Jiggins CD, Chamberlain NL, Kronforst MR, Chen R, et al. 2011. Optix drives the repeated convergent evolution of butterfly wing pattern mimicry. Science. 333(6046):1137–1141.

Reich D, Thangaraj K, Patterson N, Price AL, Singh L. 2009. Reconstructing Indian population history. Nature. 461(7263):489–494.

Roch S, Steel M. 2015. Likelihood-based tree reconstruction on a concatenation of aligned sequence data sets can be statistically inconsistent. Theor Popul Biol. 100:56–62.

Rosser N, Phillimore AB, Huertas B, Willmott KR, Mallet J. 2012. Testing historical explanations for gradients in species richness in heliconiine butterflies of tropical America. Biol J Linn Soc. 105(3):479–497.

Salazar C, Jiggins CD, Taylor JE, Kronforst MR, Linares M. 2008. Gene flow and the genealogical history of Heliconius heurippa. BMC Evol Biol. 8(1):132.

Sankararaman S, Mallick S, Dannemann M, Prüfer K, Kelso J, Pääbo S, Patterson N, Reich D. 2014. The genomic landscape of Neanderthal ancestry in present-day humans. Nature. 507(7492):354–357.

Shi CM, Yang Z. 2018. Coalescent-Based Analyses of Genomic Sequence Data Provide a Robust Resolution of Phylogenetic Relationships among Major Groups of Gibbons. Mol Biol Evol. 35(1):159–179.

Solís-Lemus C, Bastide P, Ané C. 2017. PhyloNetworks: A package for phylogenetic networks. Mol Biol Evol. 34(12):3292–3298.

Sousa V, Hey J. 2013. Understanding the origin of species with genome-scale data: Modelling gene flow. Nat Rev Genet. 14(6):404–414.

Stephens M, Smith NJ, Donnelly P. 2001. A new statistical method for haplotype reconstruction from population data. Am J Hum Genet. 68(4):978–989.

Stryjewski KF, Sorenson MD. 2017. Mosaic genome evolution in a recent and rapid avian radiation. Nat Ecol Evol. 1(12):1912–1922.

Tarasov A, Vilella AJ, Cuppen E, Nijman IJ, Prins P. 2015. Sambamba: Fast processing of NGS alignment formats. Bioinformatics. 31(12):2032–2034.

Taylor SA, Larson EL. 2019. Insights from genomes into the evolutionary importance and prevalence of hybridization in nature. Nat Ecol Evol. 3(2):170–177.

Thawornwattana Y, Dalquen D, Yang Z. 2018. Coalescent analysis of phylogenomic data confidently resolves the species relationships in the Anopheles gambiae species complex. Mol Biol Evol. 35(10):2512–2527.

Wen D, Nakhleh L. 2018. Coestimating reticulate phylogenies and gene trees from multilocus sequence data. Syst Biol. 67(3):439–457.

Xu B, Yang Z. 2016. Challenges in species tree estimation under the multispecies coalescent model. Genetics. 204(4):1353–1368.

Yang Z. 2002. Likelihood and Bayes estimation of ancestral population sizes in hominoids using data from multiple loci. Genetics. 162(4):1811–1823.

Yang Z. 2015. The BPP program for species tree estimation and species delimitation. Curr Zool. 61(5):854–865.

Yang Z, Rannala B. 2010. Bayesian species delimitation using multilocus sequence data. Proc Natl Acad Sci U S A. 107(20):9264–9269.

Yang Z, Rannala B. 2014. Unguided species delimitation using DNA sequence data from multiple loci. Mol Biol Evol. 31(12):3125–3135.

Yu Y, Dong J, Liu KJ, Nakhleh L. 2014. Maximum likelihood inference of reticulate evolutionary histories. Proc Natl Acad Sci U S A. 111(46):16448–16453.

Yu Y, Nakhleh L. 2015. A maximum pseudo-likelihood approach for phylogenetic networks. BMC Genomics. 16(Suppl 10):1–10.

Zhang C, Ogilvie HA, Drummond AJ, Stadler T. 2018. Bayesian inference of species networks from multilocus sequence data. Mol Biol Evol. 35(2):504–517.

Zhu T, Yang Z. 2012. Maximum likelihood implementation of an isolation-with-migration model with three species for testing speciation with gene flow. Mol Biol Evol. 29(10):3131–3142.

