## Supplementary figures for "Full-likelihood genomic analysis clarifies a complex history of species divergence and introgression: the example of the *erato-sara* group of *Heliconius* butterflies"

**Supplementary figures and tables**

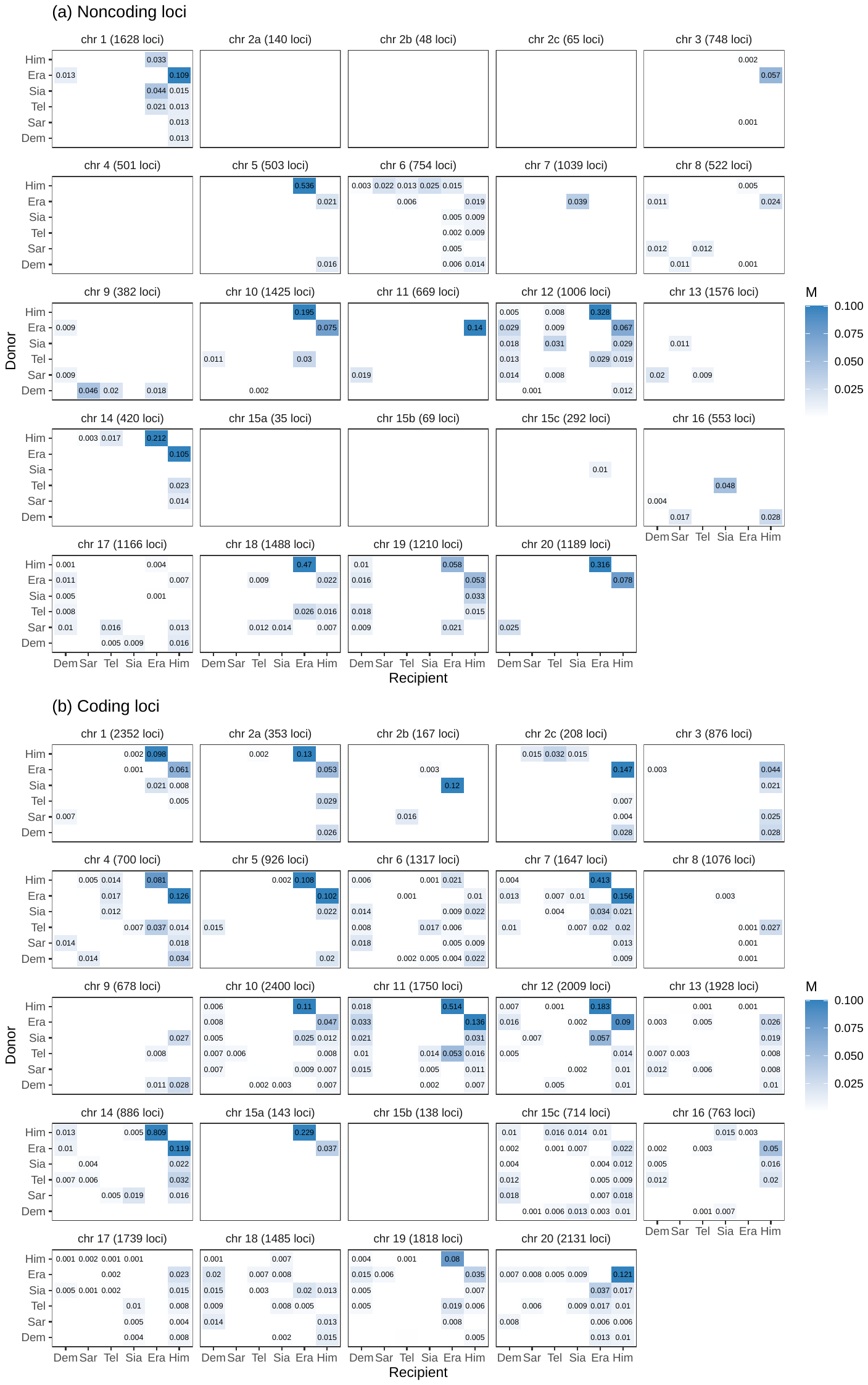
 **Figure S1.** Maximum-likelihood estimates of the pairwise migration rates (*M* = *Nm*) under the IM model from 3s for each chromosomal region in the autosomes using (**a**) noncoding and (**b**) coding loci. The donor and recipient species of gene flow are given in the y- and x-axis, respectively. *H. melpomene* was used as an outgroup. Only significant migration rate estimates (by likelihood ratio test at p < 0.01) larger than 0.001 are shown. See **Figure S2** and **Table S8** for estimates of other parameters.

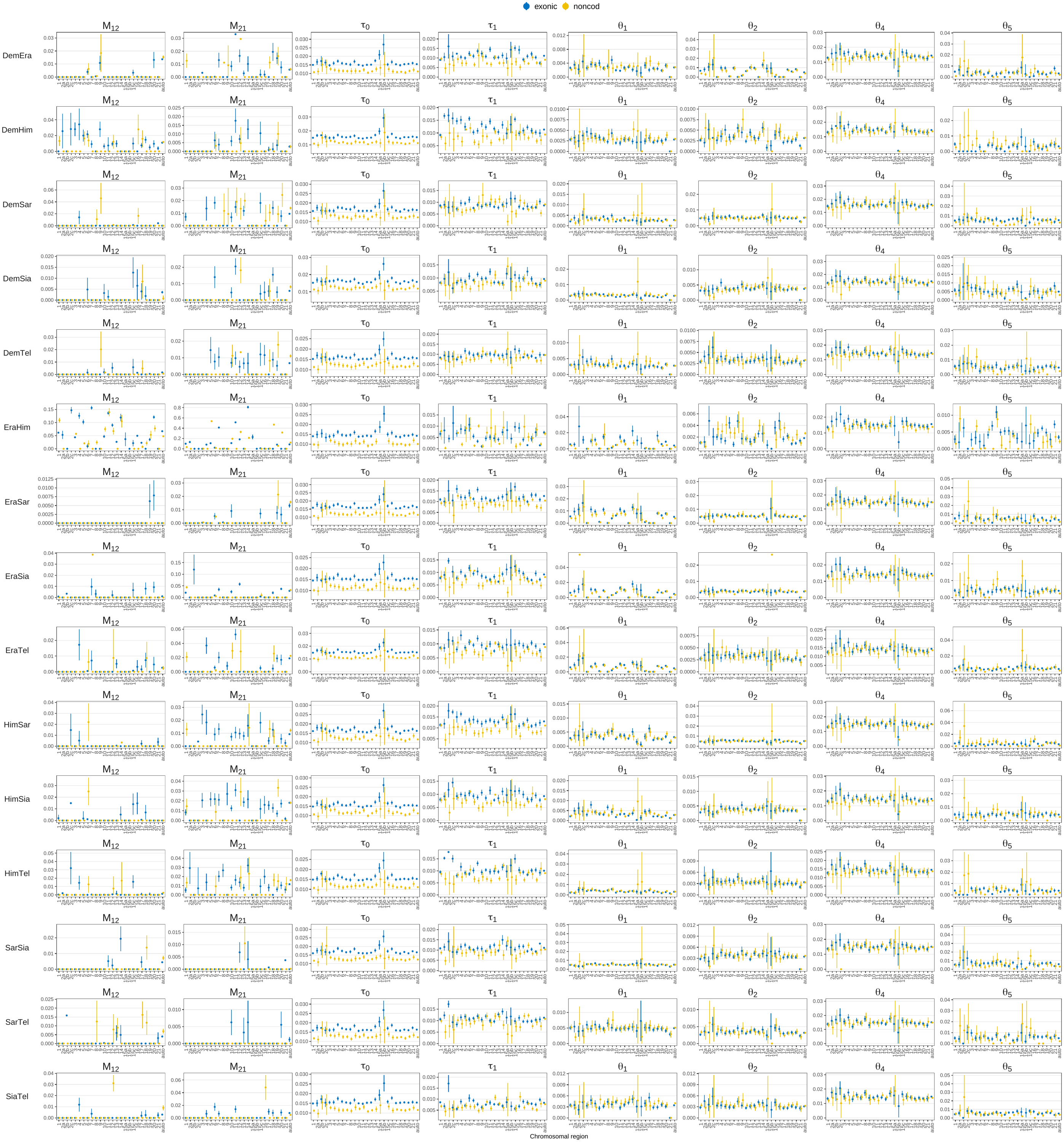

**Figure S2**. Maximum likelihood estimates of all parameters obtained in the 3s analysis under the IM model for each chromosomal region (the x-axis). The model assumes the species tree ((*S*_1_, *S*_2_), *S*_3_) and allows for gene flow between *S*_1_ and *S*_2_ at rates *M*_12_ and *M*_21_. Each locus had three sequences, with one or two sequences from *S*_1_ or *S*_2_, and at most one sequence from *S*_3_. Rows are different species pairs *S*_1_ and *S*_2_, with *H. melpomene* used as the outgroup *S*_3_ in all cases. Columns are parameters in the model. Error bars indicate two standard errors. Standard errors for some parameters for certain datasets were not shown since there were not reliably estimated. A few missing points indicate that the program did not produce reliable estimates due to numerical issues, e.g. chr 2b and 15b noncoding loci from Era/Him. See **Table S8** for complete results.

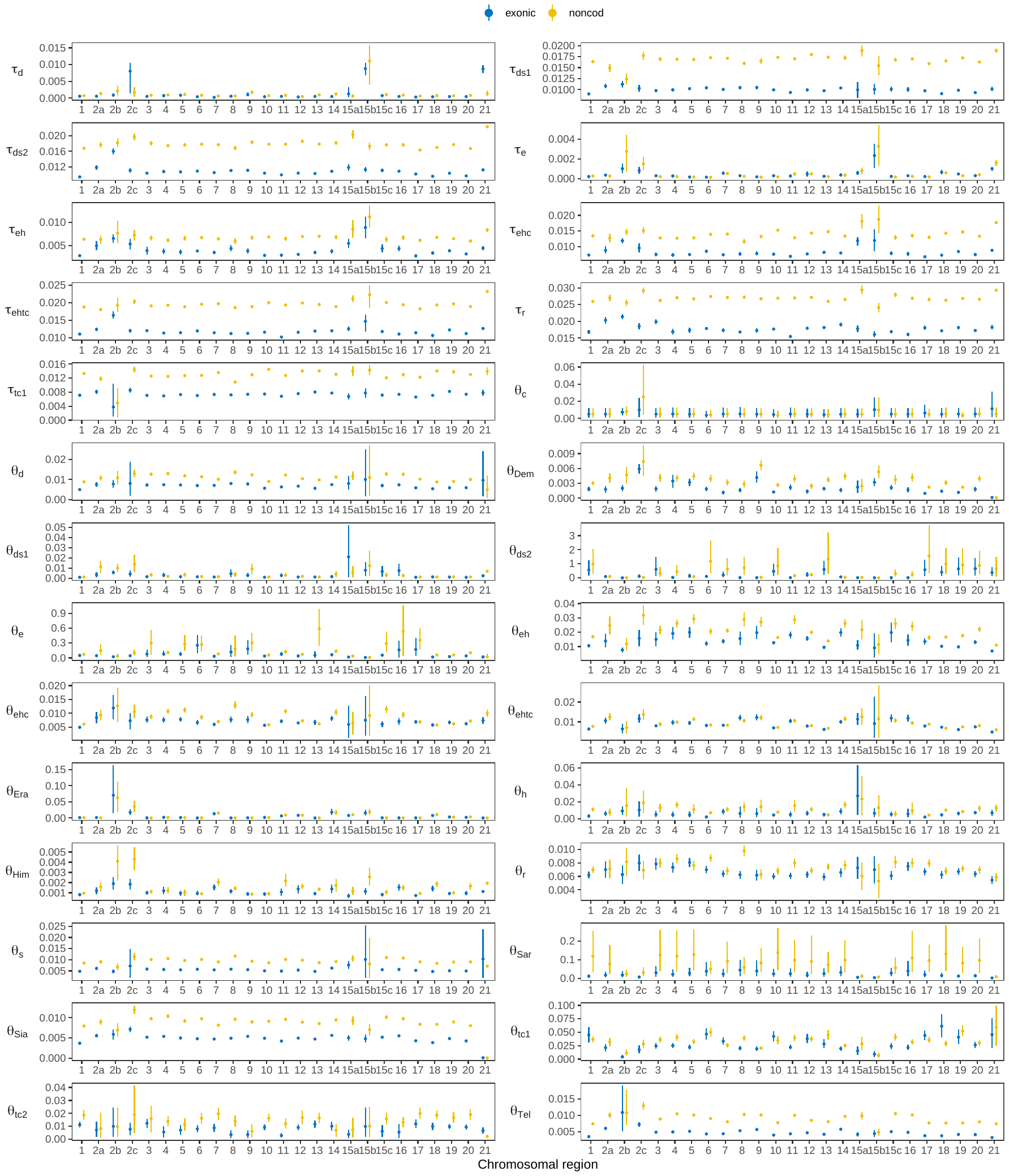

**Figure S3.** Posterior means and 95% HPD intervals of nineteen population sizes (*θ*) and nine divergence or introgression times (*τ*) under the MSci model (**Fig. 4a**) obtained using bpp for the 25 chromosomal regions (see **Table S4** for the number of loci). Only non-redundant parameters are shown; for example, *τ*_d_ ≡ *τ*_s_, so only *τ*_d_ is shown. Results for the six introgression probabilities (*ϕ*s) are in **Figure 4b**.

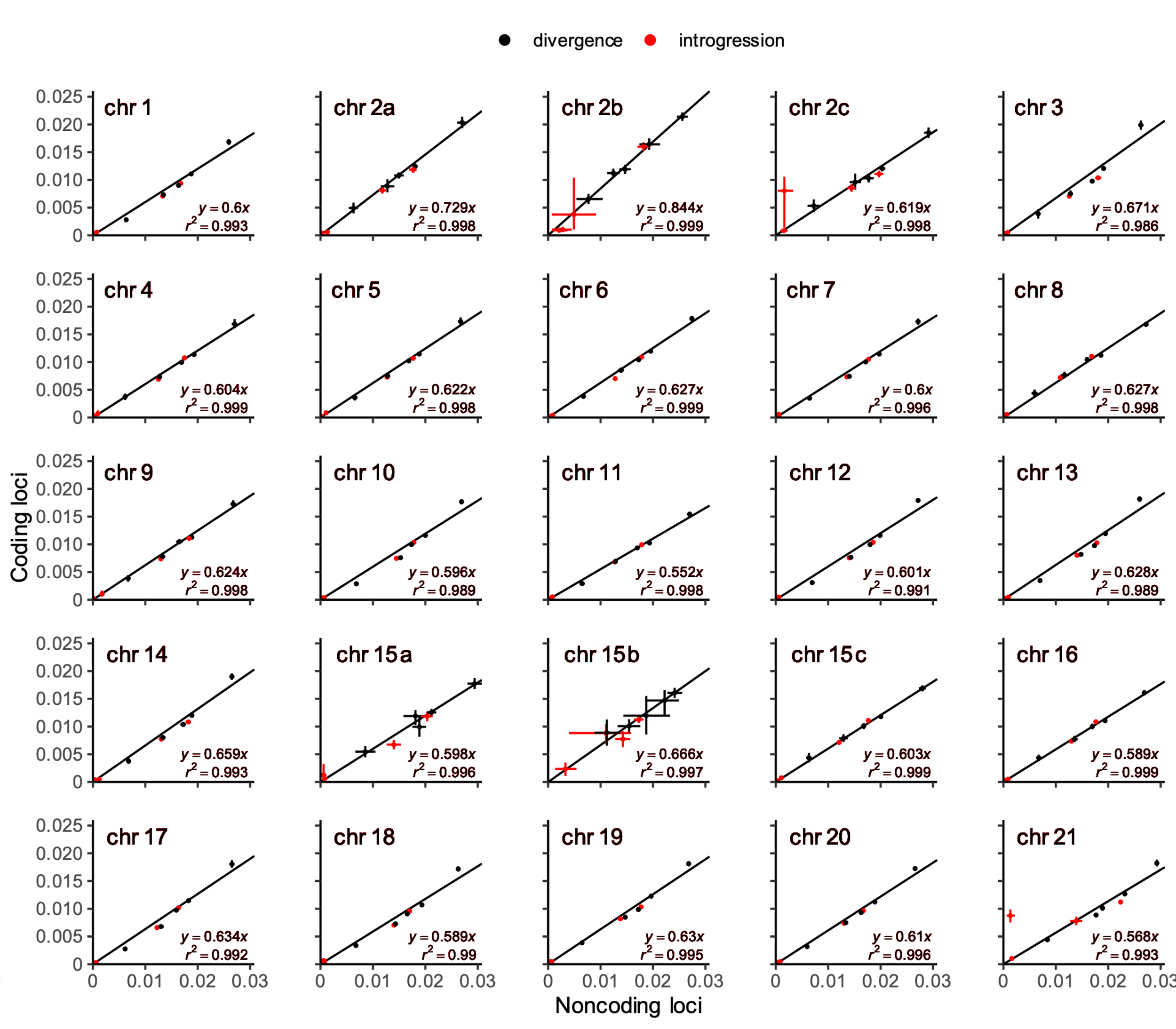

**Figure S4**. Posterior means and 95% HPD intervals of divergence times (*τ*) estimated from the noncoding (x-axis) versus coding (y-axis) loci under the MSci model of **Figure 4a**. Linear regression *y* = *c* *x* was fitted to the divergence times (black points) only, with the introgression times (red points) excluded. The outlier introgression time for chr 2c and 21 corresponds to *H. sara* → *H. demeter* introgression (*τ* _d_), which was estimated to be close to the present day for noncoding loci like other chromosomal regions, but was poorly estimated and close to the species divergence time for coding loci (**Fig. S3**).

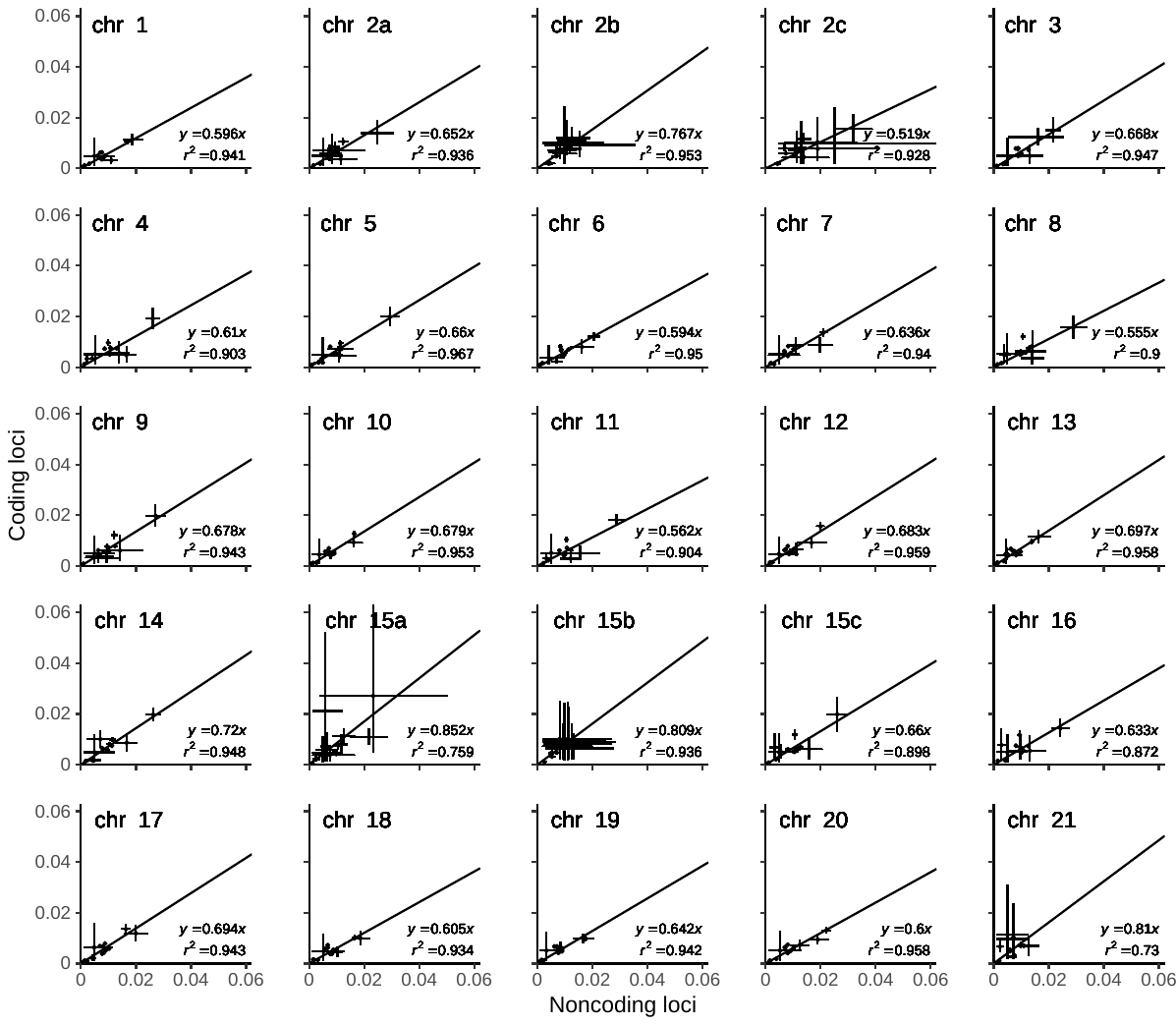

**Figure S5**. Posterior means and 95% HPD intervals of present-day and ancestral population sizes (*θ*) estimated from the noncoding (x-axis) versus coding (y-axis) loci under the MSci model (**Fig. 4a**). Linear regression *y* = *c* *x* was fitted to well-estimated population sizes, with poorly estimated *θ*_ds2_, *θ*_e_, *θ*_tc1_, *θ*_Sar_ and *θ*_Era_ excluded (see **Fig. S3**).

(a)

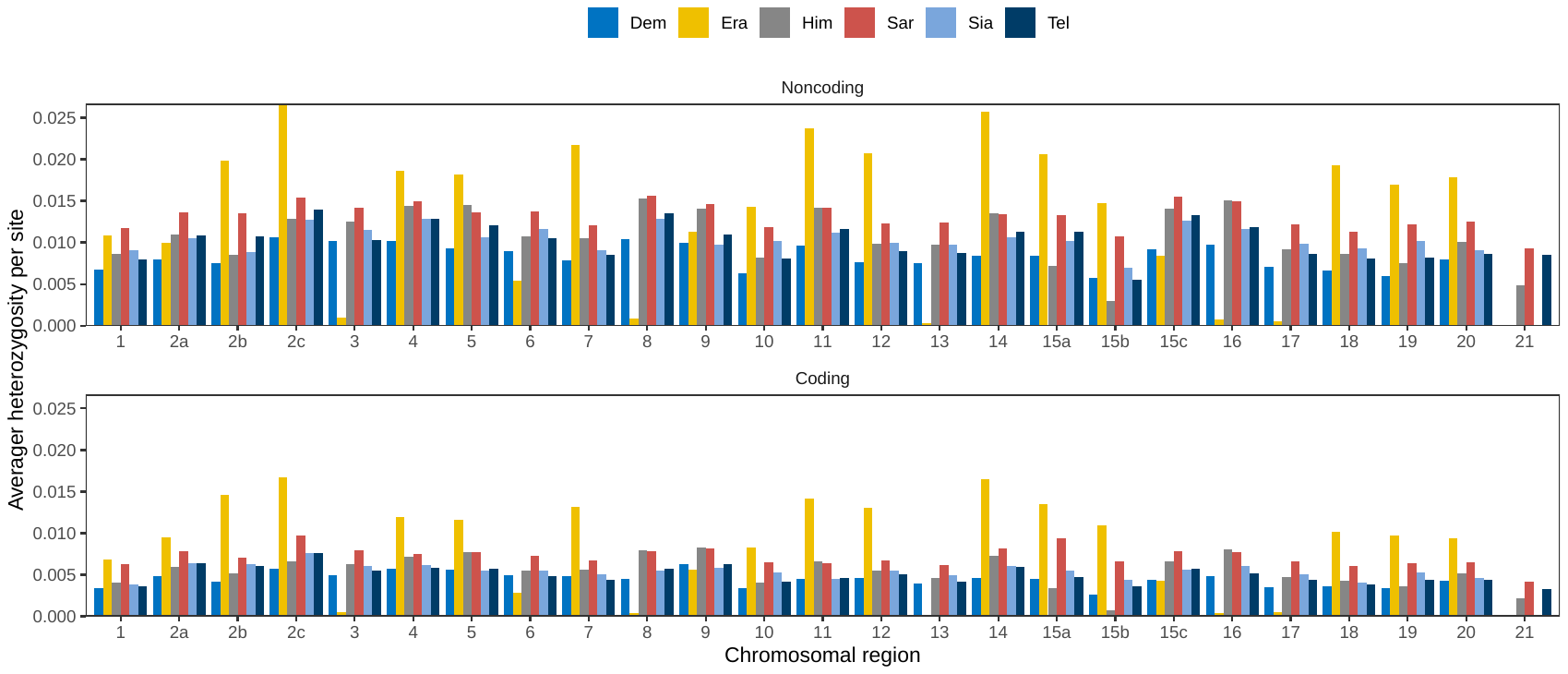

(b)

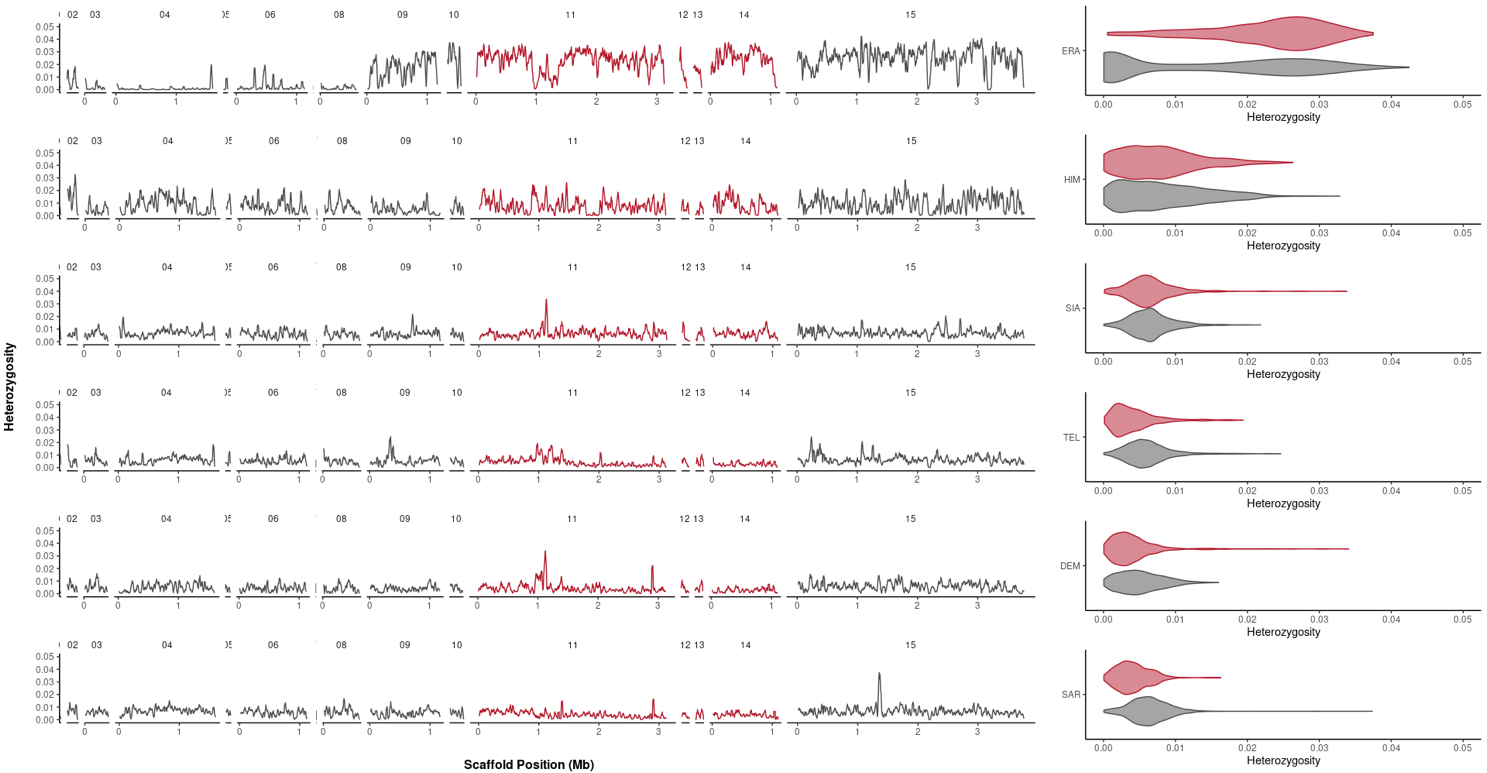

**Figure S6.** (**a**) Heterozygosity across chromosomal regions for each of the six genomes used in this study. *H. erato* individual (Era) was partially inbred and had large fluctuation of the heterozygosity level across the genome. Note that three individuals Dem, Era and Sia were females which had only one copy of the Z chromosome. (**b**) Heterozygosity in sliding windows (window size of 25 kb, step size of 5 kb) along chromosome 2 shows that the inversion region (red) is not more polymorphic than the flanking regions.

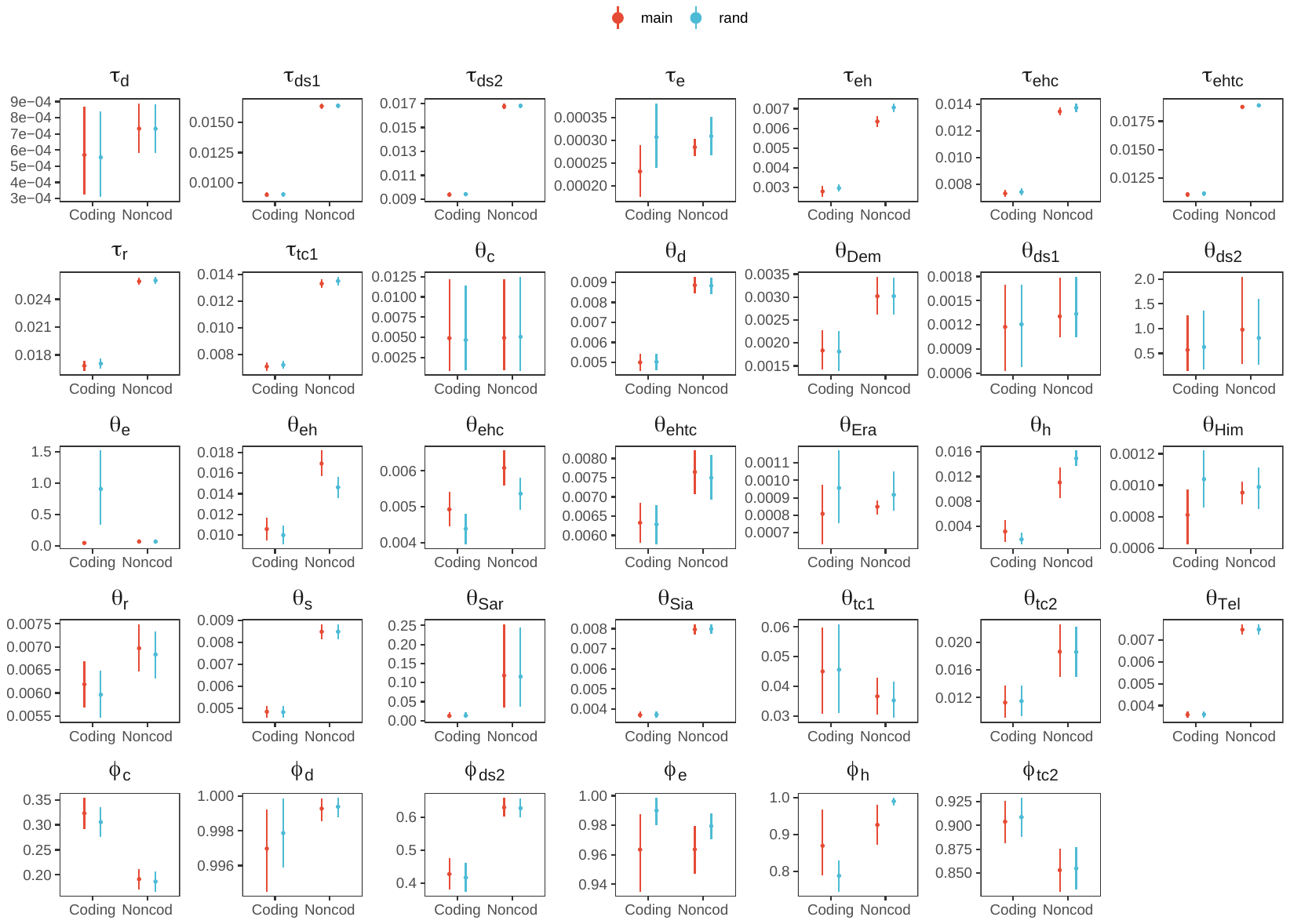

**Figure S7.** Posterior means and 95% HPD intervals of divergence/introgression times (*τ*), population sizes (*θ*) and introgression probabilities (*ϕ*) under the MSci model (**Fig. 4a**) obtained from bpp using unphased diploid sequences ('main'; **Table S9**) and randomly phased sequences ('rand') for chromosome 1 (4,902 coding loci, 6,030 noncoding loci). Only non-redundant parameters are shown; for example, *τ*_d_ ≡ *τ*_s_, so only *τ*_d_ is shown.

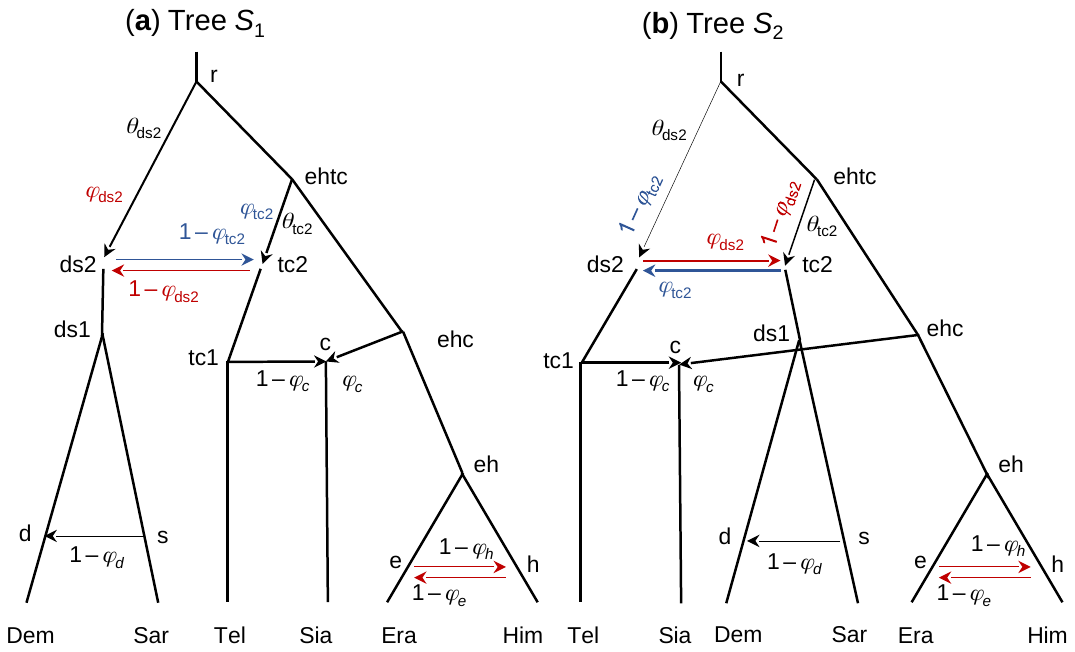

**Figure S8.** Cross-model unidentifiability of the MSci model (**Fig. 4a**). (**a**) Tree *S*_1_ is the model of **Figure 4a** used in our main analysis. (**b**) Tree *S*_2_ is another unidentifiable model with the same posterior distribution. Each of models *S*_1_ and *S*_2_ has a within-model unidentifiability due to the bidirectional introgression event at nodes e-h; see main text. Thinner lines indicate less likely paths that sequences will pass through, based on posterior estimates of introgression probabilities from bpp under the MSci model (**Fig. 4b**).

(a)

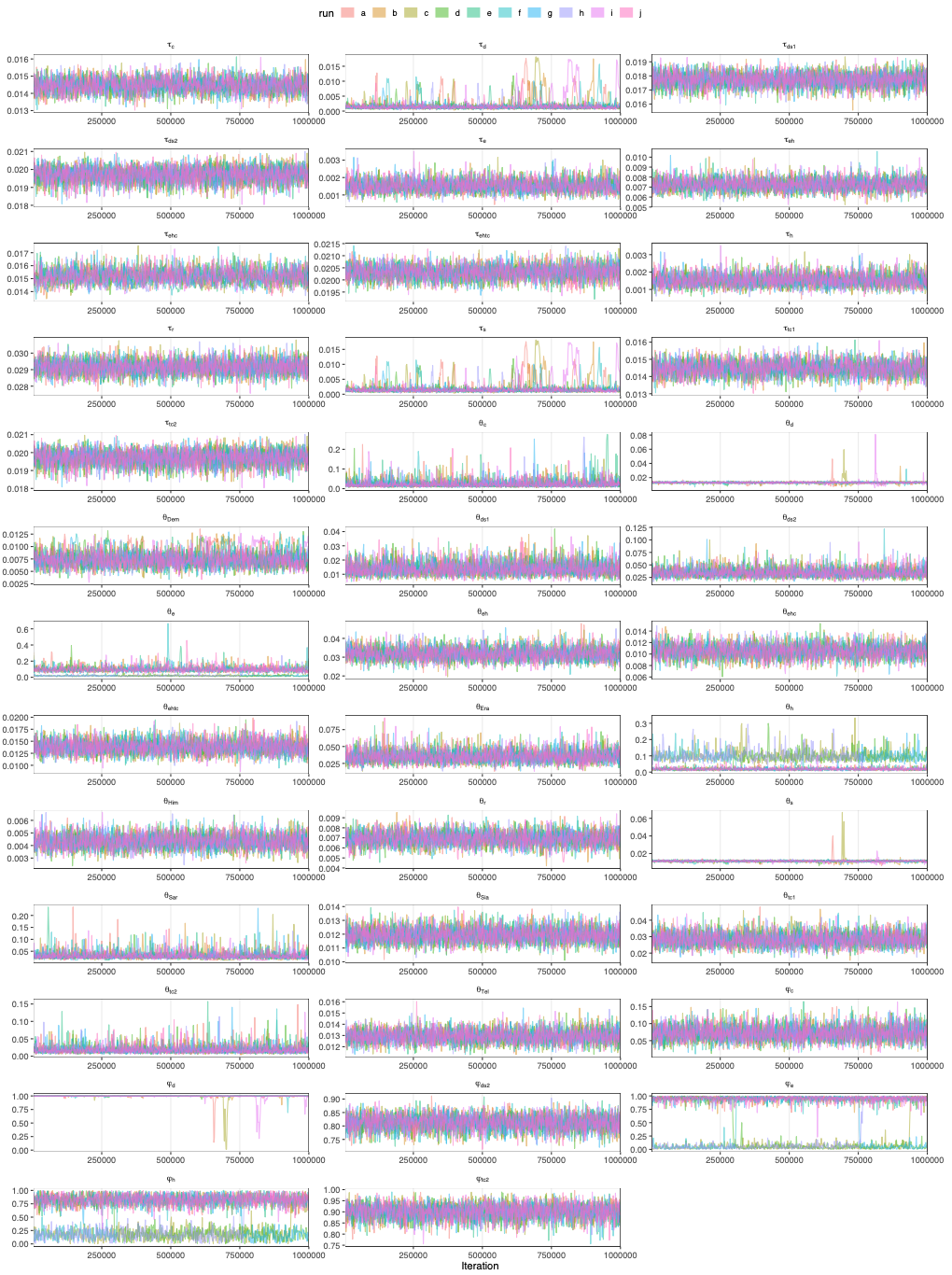

(b)

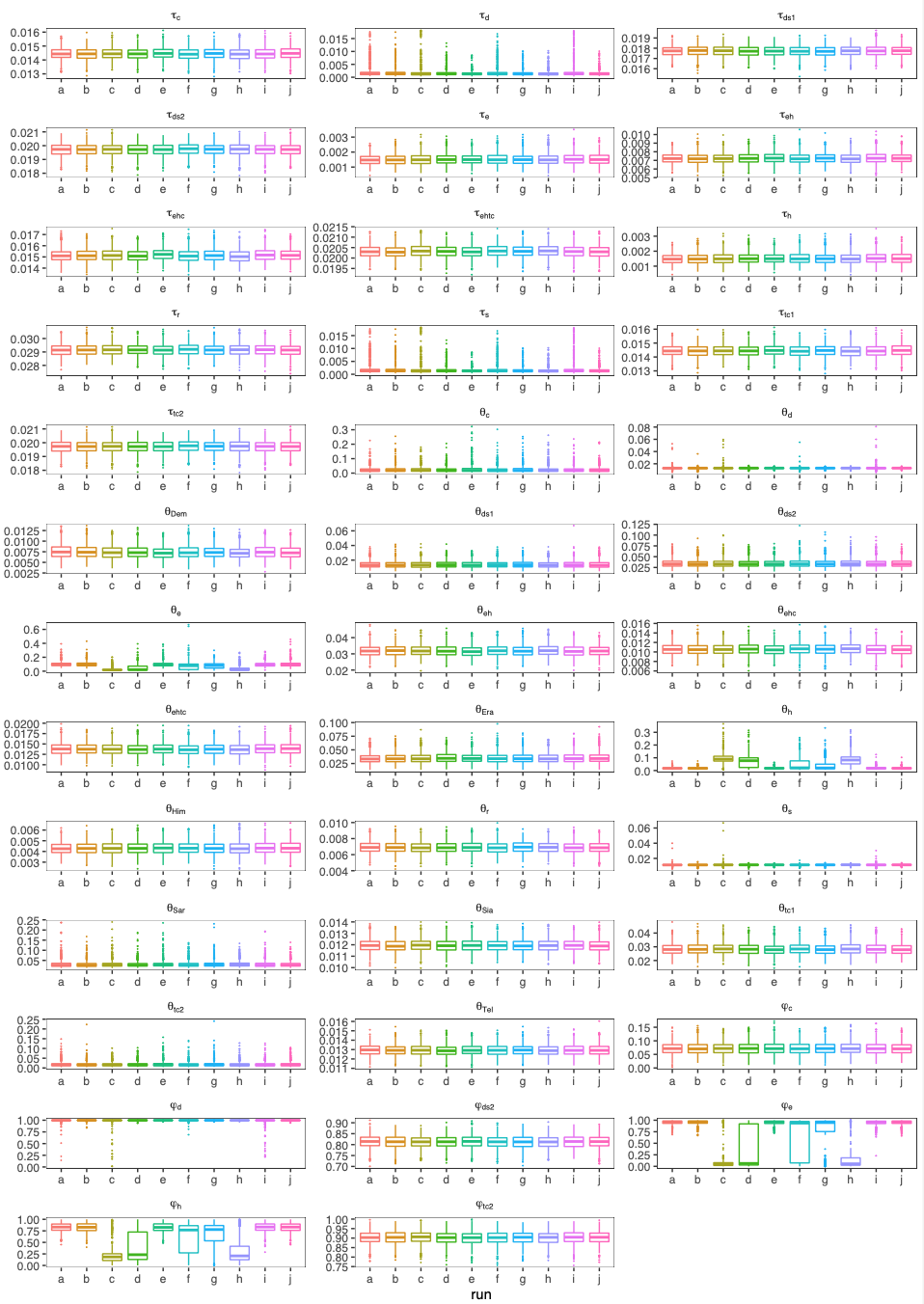

**Figure S9.** Trace plots (**a**) and box plots (**b**) of MCMC samples from bpp analysis of the MSci model (**Fig. 4a**) using 822 noncoding loci from chromosome 2c (**Table S4**). Colours represent ten independent runs. Two issues were identified in this analysis. First, runs a, c and i had mixing issues (for parameters *ϕ*_d_, *θ*_10d_ and *θ*_11s_) and those runs were discarded. Second, there were label-switching unidentifiability issues for the Era–Him introgression and thus the samples were pre-processed before summarizing: if *ϕ*_e_ < 0.5, we changed *ϕ*_e_ to 1 – *ϕ*_e_, and *ϕ*_h_ to 1 – *ϕ*_h_, and swapped *θ*_e_ and *θ*_h_ (Flouri et al., 2020).

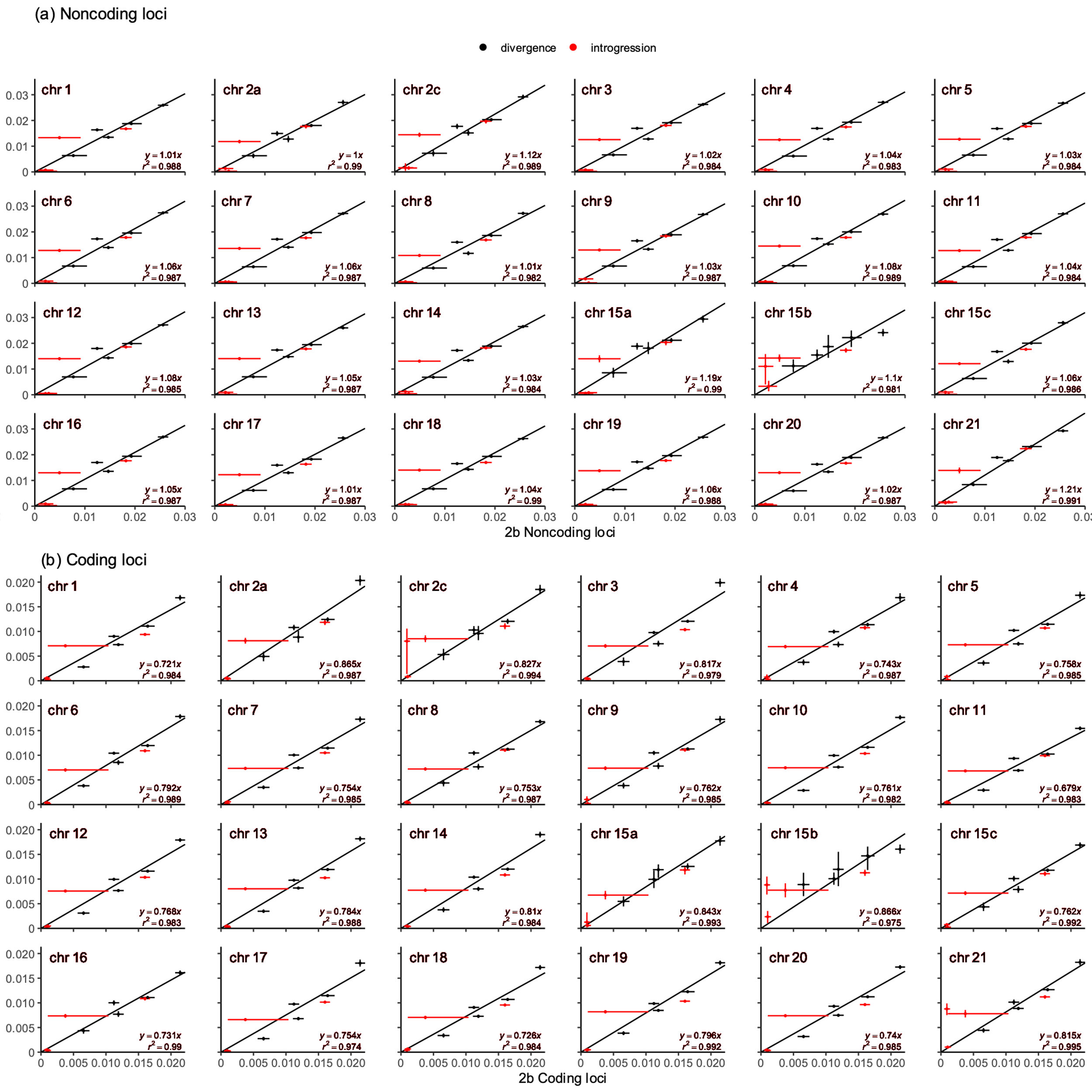

**Figure S10**. Posterior means and 95% HPD intervals of divergence times (*τ*) estimated from the chromosome 2b inversion region and other chromosomal regions for (**a**) noncoding and (**b**) coding loci. The slope *c* of the regression *y* = *c* *x* measures the relative mutation rate of other region of the genome (*y*) relative to the 2b region (*x*). The long horizontal bar in each plot corresponds to an uncertain estimate of the introgression time from *H. telesiphe* into *H. hecalesia* (*τ*_tc1_), indicating the absence of such introgression in the 2b region. The divergence time between *H. demeter* and *H. sara* (*τ*_ds1_) is always above the regression line, corresponding to the younger estimate of *τ*_ds1_ in the chromosome 2b region compared to other chromosomes (see **Fig. S3**).

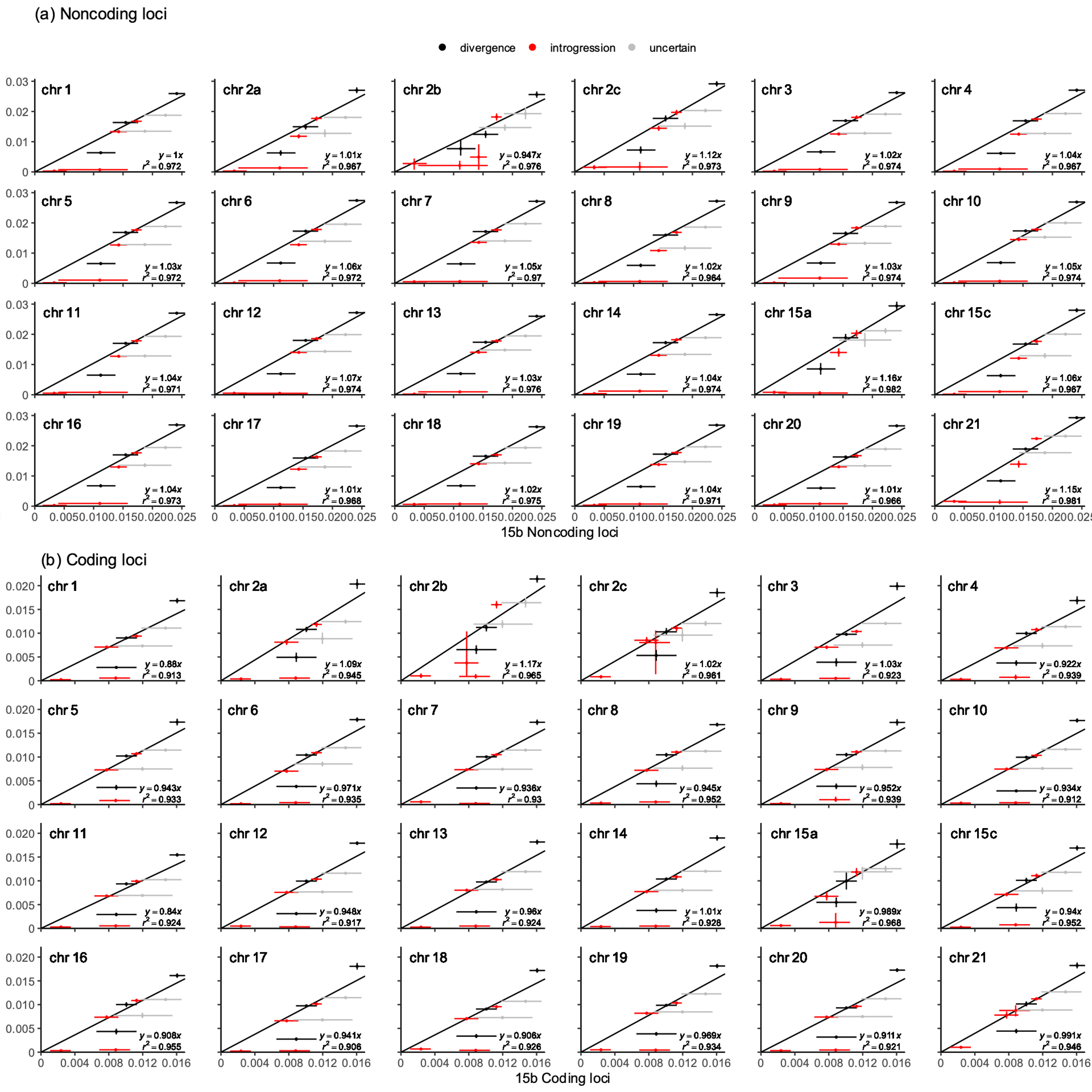

**Figure S11**. Posterior means and 95% HPD intervals of divergence times (*τ*) estimated from the chromosome 15b inversion region and other chromosomal regions for (**a**) noncoding and (**b**) coding loci. The slope *c* of the regression *y* = *c* *x* measures the relative mutation rate of other region of the genome (*y*) relative to the 15b region (*x*). The grey points correspond to *τ*_ehtc_ and *τ*_ehc_ that were poorly estimated (see **Fig. 6b**, Scenario 1) and were excluded from the regression fitting. The time *τ*_d_ for *H. sara* → *H. demeter* introgression was estimated to be close to the divergence time in 15b, but with high uncertainty.

**Table S1. Sample information.**

| **Clade** | **Species** | **Subspecies** | **Abbreviation** | **Sex** | **BROAD ID** | **Mean depth** | | **Dataset** | |
| --- | --- | --- | --- | --- | --- | --- | --- | --- | --- |
|  |  |  |  |  |  | **hmelv25 ref** | **heradem ref** | **bpp** | **3s** |
| *erato* | *H. erato* | *demophoon* | Era | F | SM-6OSJR | 37.5 | 66.6 | yes | yes |
|  | *H. himera* | *-* | Him | M | SM-6OSJQ | 22.0 | 31.5 | yes | yes |
|  | *H. hecalesia* | *formosus* | Sia | F | SM-ADRBM | 53.2 | 71.8 | yes | yes |
|  | *H. telesiphe* | *telesiphe* | Tel | M | SM-ADRBQ | 22.0 | 28.2 | yes | yes |
| *sara* | *H. demeter* | *-* | Dem | F | SM-6OSKP | 40.5 | 50.4 | yes | yes |
|  | *H. sara* | *magdalena* | Sar | M | SM-ADRBN | 47.5 | 57.7 | yes | yes |
| *melpomene* | *H. melpomene* | *melpomene* | Mel | F | SM-5BSJX | 104.2 | 48.7 | no | yes |

**Table S2.** Estimates of the base-calling error rate for homozygous reference allele (GT00) and homozygous alternative allele (GT11).

| **minDP** | **Genotype** | **Dem** | **Era** | **Him** | **Sar** | **Sia** | **Tel** | **Mel** |
| --- | --- | --- | --- | --- | --- | --- | --- | --- |
| 20 | GT00 | 0.06% | 0.08% | 0.11% | 0.06% | 0.06% | 0.10% | 0.08% |
|  | GT11 | 0.30% | 0.33% | 0.29% | 0.21% | 0.22% | 0.27% | 0.20% |
| 50 | GT00 | 0.07% | 0.08% | 0.53% | 0.06% | 0.06% | 0.30% | 0.08% |
|  | GT11 | 0.29% | 0.32% | 1.33% | 0.20% | 0.21% | 0.72% | 0.18% |

**Table S3.** Estimates of the base-calling error rate for homozygous reference allele (GT00) and homozygous alternative allele (GT11) by genomic region (coding and noncoding) using different depth (DP) filters: (1) Dpdef: minDP ≥ 12, (2) Dpmin: minDP ≥ 0.5*meanDP, (3) Dpmax: maxDP ≤ 2*meanDP, (4) Dpbot: minDP ≥ 0.5*meanDP and maxDP ≤ 2*meanDP.

| **Genotype** | **Region** | **Filter** | **Dem** | **Era** | **Him** | **Sar** | **Sia** | **Tel** | **Mel** |
| --- | --- | --- | --- | --- | --- | --- | --- | --- | --- |
| GT00 | noncoding | Dpdef | 0.07% | 0.08% | 0.10% | 0.06% | 0.06% | 0.12% | 0.07% |
|  |  | Dpmin | 0.07% | 0.08% | 0.10% | 0.06% | 0.06% | 0.12% | 0.07% |
|  |  | Dpmax | 0.05% | 0.05% | 0.07% | 0.05% | 0.05% | 0.06% | 0.07% |
|  |  | Dpbot | 0.05% | 0.05% | 0.07% | 0.05% | 0.05% | 0.06% | 0.07% |
|  | coding | Dpdef | 0.06% | 0.07% | 0.09% | 0.05% | 0.06% | 0.08% | 0.08% |
|  |  | Dpmin | 0.06% | 0.07% | 0.09% | 0.05% | 0.06% | 0.08% | 0.08% |
|  |  | Dpmax | 0.05% | 0.06% | 0.08% | 0.05% | 0.05% | 0.05% | 0.08% |
|  |  | Dpbot | 0.05% | 0.06% | 0.08% | 0.05% | 0.05% | 0.05% | 0.08% |
| GT11 | noncoding | Dpdef | 0.28% | 0.29% | 0.21% | 0.19% | 0.20% | 0.22% | 0.12% |
|  |  | Dpmin | 0.26% | 0.28% | 0.21% | 0.18% | 0.19% | 0.22% | 0.10% |
|  |  | Dpmax | 0.26% | 0.27% | 0.16% | 0.18% | 0.18% | 0.16% | 0.11% |
|  |  | Dpbot | 0.24% | 0.25% | 0.16% | 0.18% | 0.17% | 0.16% | 0.10% |
|  | coding | Dpdef | 0.17% | 0.20% | 0.13% | 0.10% | 0.11% | 0.14% | 0.16% |
|  |  | Dpmin | 0.17% | 0.20% | 0.13% | 0.10% | 0.11% | 0.14% | 0.15% |
|  |  | Dpmax | 0.16% | 0.17% | 0.10% | 0.09% | 0.09% | 0.09% | 0.13% |
|  |  | Dpbot | 0.15% | 0.17% | 0.10% | 0.09% | 0.09% | 0.09% | 0.12% |

**Table S4.** Number of loci for each of the 25 chromosomal regions in the bpp dataset (in blocks of 100 loci or 200 loci) and in the 3s dataset.

| **Chr** | **bpp dataset (6 genomes)** | | | | | | | | | | | | | | **3s dataset**  **(7 genomes)** | |
| --- | --- | --- | --- | --- | --- | --- | --- | --- | --- | --- | --- | --- | --- | --- | --- | --- |
|  | **Noncoding loci** | | | | | | | **Coding loci** | | | | | | | **#noncoding loci** | **#coding loci** |
|  | **#loci** | **100-locus dataset** | | | **200-locus dataset** | | | **#loci** | **100-locus dataset** | | | **200-locus dataset** | | |  |  |
|  |  | **#blocks** | **#loci in last block** | **last block discarded** | **#blocks** | **#loci in last block** | **last block discarded** |  | **#blocks** | **#loci in last block** | **last block discarded** | **#blocks** | **#loci in last block** | **last block discarded** |  |  |
| **1** | 6030 | 61 | 30 | yes | 31 | 30 | yes | 4902 | 50 | 2 | yes | 25 | 102 | no | 1628 | 2352 |
| **2a** | 1275 | 13 | 75 | no | 7 | 75 | no | 1263 | 13 | 63 | no | 7 | 63 | no | 140 | 353 |
| **2b** | 411 | 5 | 11 | yes | 3 | 11 | yes | 516 | 6 | 16 | yes | 3 | 116 | no | 48 | 167 |
| **2c** | 822 | 9 | 22 | yes | 5 | 22 | yes | 772 | 8 | 72 | no | 4 | 172 | no | 65 | 208 |
| **3** | 3621 | 37 | 21 | yes | 19 | 21 | yes | 2410 | 25 | 10 | yes | 13 | 10 | yes | 748 | 876 |
| **4** | 3351 | 34 | 51 | no | 17 | 151 | no | 2025 | 21 | 25 | yes | 11 | 25 | yes | 501 | 700 |
| **5** | 3376 | 34 | 76 | no | 17 | 176 | no | 2389 | 24 | 89 | no | 12 | 189 | no | 503 | 926 |
| **6** | 4674 | 47 | 74 | no | 24 | 74 | no | 3932 | 40 | 32 | yes | 20 | 132 | no | 754 | 1317 |
| **7** | 5060 | 51 | 60 | no | 26 | 60 | no | 3935 | 40 | 35 | yes | 20 | 135 | no | 1039 | 1647 |
| **8** | 3147 | 32 | 47 | no | 16 | 147 | no | 2829 | 29 | 29 | yes | 15 | 29 | yes | 522 | 1076 |
| **9** | 2870 | 29 | 70 | no | 15 | 70 | no | 1828 | 19 | 28 | yes | 10 | 28 | yes | 382 | 678 |
| **10** | 6462 | 65 | 62 | no | 33 | 62 | no | 5519 | 56 | 19 | yes | 28 | 119 | no | 1425 | 2400 |
| **11** | 3964 | 40 | 64 | no | 20 | 164 | no | 4090 | 41 | 90 | no | 21 | 90 | no | 669 | 1750 |
| **12** | 5602 | 57 | 2 | yes | 29 | 2 | yes | 5312 | 54 | 12 | yes | 27 | 112 | no | 1006 | 2009 |
| **13** | 6186 | 62 | 86 | no | 31 | 186 | no | 4471 | 45 | 71 | no | 23 | 71 | no | 1576 | 1928 |
| **14** | 3062 | 31 | 62 | no | 16 | 62 | no | 2580 | 26 | 80 | no | 13 | 180 | no | 420 | 886 |
| **15a** | 491 | 5 | 91 | no | 3 | 91 | no | 493 | 5 | 93 | no | 3 | 93 | no | 35 | 143 |
| **15b** | 149 | 2 | 49 | no | 1 | 149 | no | 167 | 2 | 67 | no | 1 | 167 | no | 69 | 138 |
| **15c** | 2832 | 29 | 32 | yes | 15 | 32 | yes | 2228 | 23 | 28 | yes | 12 | 28 | yes | 292 | 714 |
| **16** | 3479 | 35 | 79 | no | 18 | 79 | no | 2090 | 21 | 90 | no | 11 | 90 | no | 553 | 763 |
| **17** | 5017 | 51 | 17 | yes | 26 | 17 | yes | 4531 | 46 | 31 | yes | 23 | 131 | no | 1166 | 1739 |
| **18** | 5914 | 60 | 14 | yes | 30 | 114 | no | 3692 | 37 | 92 | no | 19 | 92 | no | 1488 | 1485 |
| **19** | 5734 | 58 | 34 | yes | 29 | 134 | no | 4533 | 46 | 33 | yes | 23 | 133 | no | 1210 | 1818 |
| **20** | 5087 | 51 | 87 | no | 26 | 87 | no | 4864 | 49 | 64 | no | 25 | 64 | no | 1189 | 2131 |
| **21** | 4350 | 44 | 50 | no | 22 | 150 | no | 3628 | 37 | 28 | yes | 19 | 28 | yes | 574 | 1348 |
| Total | 92966 | 933 | - | - | 472 | - | - | 74999 | 749 | - | - | 382 | - | - | 18002 | 29552 |

**Table S5.** Proportions of species tree estimates (MAP trees, with minimum, median and maximum posterior probabilities shown in parentheses) from the blockwise bpp MSC analysis using blocks of 100 loci and 200 loci, summarized into four major regions: auto (all autosomes excluding 2b and 15b inversions), 2b inversion, 15b inversion and the Z chromosome (chr 21). n is the number of blocks. Trees are ordered in decreasing proportions of blocks for the 100-locus dataset. Tree indices correspond to those of **Figure 2**.

| **Region** | **Group** | **Tree** | **Tree index** | **100-locus dataset** | | **200-locus dataset** | |
| --- | --- | --- | --- | --- | --- | --- | --- |
|  |  |  |  | **n** | **Proportion** | **n** | **Proportion** |
| **Noncoding** | auto | ((Dem, Sar), ((Era, Him), (Sia, Tel))); | i | 511 | 0.579 (0.39, 1.00, 1.00) | 278 | 0.622 (0.44, 1.00, 1.00) |
|  |  | ((Dem, Sar), (((Era, Him), Sia), Tel)); | ii | 356 | 0.403 (0.48, 1.00, 1.00) | 168 | 0.376 (0.50, 1.00, 1.00) |
|  |  | ((Dem,  Sar), (((Era, Him), Tel), Sia)); | vi | 11 | 0.013 (0.44, 0.69, 0.95) | 1 | 0.002 (0.50, 0.50, 0.50) |
|  |  | (((Dem, Sar), (Sia, Tel)), (Era, Him)); | iv | 5 | 0.006 (0.87, 0.95, 1.00) | 0 | 0 |
|  | 2b | (((Dem, Sar), Tel), ((Era, Him), Sia)); | iii | 4 | 1.000 (1.00, 1.00, 1.00) | 2 | 1.000 (1.00, 1.00, 1.00) |
|  | 15b | (((Dem, Sar), (Sia, Tel)), (Era, Him)); | iv | 2 | 1.000 (0.66, 0.83, 1.00) | 1 | 1.000 (1.00, 1.00, 1.00) |
|  | 21 | ((Dem, Sar), (((Era, Him), Sia), Tel)); | ii | 43 | 0.977 (0.92, 1.00, 1.00) | 22 | 1.000 (1.00, 1.00, 1.00) |
|  |  | (((Dem, Sar), Tel), ((Era, Him), Sia)); | iii | 1 | 0.023 (0.98, 0.98, 0.98) | 0 | 0 |
| **Coding** | auto | ((Dem, Sar), ((Era, Him), (Sia, Tel))); | i | 331 | 0.469 (0.43, 0.99, 1.00) | 179 | 0.497 (0.55, 1.00, 1.00) |
|  |  | ((Dem, Sar), (((Era, Him), Sia), Tel)); | ii | 327 | 0.463 (0.42, 0.98, 1.00) | 171 | 0.475 (0.52, 1.00, 1.00) |
|  |  | (((Dem, Sar), (Sia, Tel)), (Era, Him)); | iv | 21 | 0.030 (0.47, 0.75, 1.00) | 6 | 0.017 (0.53, 0.98, 1.00) |
|  |  | ((Dem, Sar), (((Era, Him), Tel), Sia)); | vi | 13 | 0.018 (0.40, 0.61, 0.96) | 3 | 0.008 (0.37, 0.38, 0.86) |
|  |  | (((Dem, Sar), Tel), ((Era, Him), Sia)); | iii | 5 | 0.007 (0.59, 0.68, 0.94) | 0 | 0 |
|  |  | (((Dem, Sar), ((Era, Him), Sia)), Tel); | viii | 3 | 0.004 (0.46, 0.49, 0.68) | 0 | 0 |
|  |  | ((Dem, ((Era, Him), (Sia, Tel))), Sar); | x | 3 | 0.004 (0.48, 0.53, 0.85) | 0 | 0 |
|  |  | (((Dem, ((Era, Him), Sia)), Tel), Sar); | v | 1 | 0.001 (0.95, 0.95, 0.95) | 1 | 0.003 (1.00, 1.00, 1.00) |
|  |  | (((Dem, Sar), Sia), ((Era, Him), Tel)); | vii | 1 | 0.001 (0.69, 0.69, 0.69) | 0 | 0 |
|  |  | ((Dem, (((Era, Him), Sia), Tel)), Sar); | ix | 1 | 0.001 (0.90, 0.90, 0.90) | 0 | 0 |
|  | 2b | (((Dem, Sar), Tel), ((Era, Him), Sia)); | iii | 5 | 1.000 (1.00, 1.00, 1.00) | 3 | 1.000 (1.00, 1.00, 1.00) |
|  | 15b | (((Dem, Sar), (Sia, Tel)), (Era, Him)); | iv | 2 | 1.000 (0.84, 0.92, 1.00) | 1 | 1.000 (1.00, 1.00, 1.00) |
|  | 21 | ((Dem, Sar), (((Era, Him), Sia), Tel)); | ii | 35 | 0.972 (0.85, 1.00, 1.00) | 18 | 1.000 (0.99, 1.00, 1.00) |
|  |  | (((Dem, Sar), Tel), ((Era, Him), Sia)); | iii | 1 | 0.028 (0.56, 0.56, 0.56) | 0 | 0 |

**Table S6.** Proportions of species tree estimates (MAP trees), with minimum (pr_min), median (pr_med) and maximum (pr_max) posterior probabilities, from the bpp MSC analysis using blocks of 100 loci, summarized by chromosomal region.

*See excel file.*

**Table S7.** Proportions of species tree estimates (MAP trees), with minimum (pr_min), median (pr_med) and maximum (pr_max) posterior probabilities, from the bpp MSC analysis using blocks of 200 loci, summarized by chromosomal region.

*See excel file.*

**Table S8.** Maximum likelihood estimates (MLEs) of parameters under the isolation-with-migration (IM) model obtained from 3s for each of the fifteen pairs of species. LRT (likelihood ratio test) statistic was used to select the best-fit model with or without gene flow (M2 vs M0), using the p-value threshold of 1%. See **Figure S2** for plots of these estimates. ‘auto’ used all autosomal loci excluding the two inversion regions (see **Table S4** for the number of loci).

*See excel file.*

**Table S9.** Posterior means and 95% HPD intervals (in parentheses) of parameters under the MSci model in **Figure 4a** obtained from bpp using noncoding and coding loci from each chromosomal region.

*See excel file.*
